# Definition of germ layer cell lineage alternative splicing programs reveals a critical role for Quaking in specifying cardiac cell fate

**DOI:** 10.1101/2020.12.22.423880

**Authors:** W. Samuel Fagg, Naiyou Liu, Ulrich Braunschweig, Karen Larissa Pereira de Castro, Xiaoting Chen, Frederick S. Ditmars, Steven G. Widen, John Paul Donohue, Katalin Modis, William K. Russell, Jeffrey H. Fair, Matthew T. Weirauch, Benjamin J. Blencowe, Mariano A. Garcia-Blanco

## Abstract

Alternative splicing is critical for development; however, its role in the specification of the three embryonic germ layers is poorly understood. By performing RNA-Seq on human embryonic stem cells (hESCs) and derived definitive endoderm, cardiac mesoderm, and ectoderm cell lineages, we detect distinct alternative splicing programs associated with each lineage. The most prominent splicing program differences are observed between definitive endoderm and cardiac mesoderm. Integrative multi-omics analyses link each program with lineage-enriched RNA binding protein regulators, and further suggest a widespread role for Quaking (QKI) in the specification of cardiac mesoderm. Remarkably, knockout of QKI disrupts the cardiac mesoderm-associated alternative splicing program and formation of myocytes. These changes arise in part through reduced expression of *BIN1* splice variants linked to cardiac development. Mechanistically, we find that QKI represses inclusion of exon 7 in *BIN1* pre-mRNA via an exonic ACUAA motif, and this is concomitant with intron removal and cleavage from chromatin. Collectively, our results uncover alternative splicing programs associated with the three germ lineages and demonstrate an important role for QKI in the formation of cardiac mesoderm.

## INTRODUCTION

Pluripotent stem cells (PSCs), such as embryonic stem cells (ESCs) and induced pluripotent stem cells, can differentiate into any adult somatic cell (1). They are also valuable models for the study of development (2), regenerative medicine applications (3, 4), and drug discovery/toxicity testing (5). Recent advances in human PSC differentiation have enabled the production of large numbers of nearly pure germ layer committed progenitors including definitive endoderm, mesoderm, ectoderm, and their progeny (6–10). This has enabled the detailed characterization of the transcriptional and epigenetic/chromatin states of these lineages in comparison with hPSCs, underscoring the diversity of their transcriptional outputs (7,8,11–14). However, the use of these approaches to investigate post-transcriptional regulatory programs underlying germ layer specification remains largely unexplored.

Alternative splicing enables the widespread quantitative and qualitative regulation of gene outputs (15, 16). Accordingly, different cell and tissue types have unique splicing patterns that contribute to their identity, such as those observed in muscle and brain (17, 18). Studies have also identified splicing patterns that are unique to stem and precursor cells, including embryonic stem cells (ESCs) compared to cardiac precursor cells (19, 20), other late mesoderm cell types (21), and neural stem cells (22, 23). Tissue- and developmental stage-specific alternative splicing, and indeed most RNA processing, are controlled in large part by RNA binding proteins (RBPs) (24, 25).

Quaking (QKI) proteins represent one such family of RBPs. From the single *Quaking* locus in human and mouse at least three major RNA and protein isoforms are produced (26, 27). Like other RBPs, QKI is multifunctional, regulating both nuclear and cytoplasmic RNA processing (28–32), with the QKI5 isoform being necessary and sufficient for splicing function (33). Although there is a clear role for *QKI* in the transition from myoblasts to myotubes (29), in smooth muscle (34, 35) and cardiac muscle (36), it is not known whether QKI regulates earlier cell fate decisions. Other RBPs, including RBFOX2, hnRNPF/H, and MBNL family members, are known to regulate alternative splicing programs that control pluripotency, self-renewal, and differentiation (22,37,38). However, the extent to which splicing programs and regulatory RBPs diverge during early cell fate lineage commitment, and their influence on downstream differentiation programs is unknown.

Here we used RNA-sequencing (RNA-seq) to address these questions by quantitatively profiling both developmentally-regulated and lineage-specific alternative splicing, as well as RBP abundance dynamics, in H9 hESCs differentiated in parallel to definitive endoderm (DE), cardiac mesoderm (CM), or neuroectoderm (ECT) cells. These analyses provide evidence for distinct lineage-specific splicing programs, with a particularly marked difference between DE and CM cells. In fact, many alternative exons are spliced in an opposite manner in DE and CM cells. We further show that many of these differentially spliced exons contain – or are flanked by – motifs that are bound and regulated by QKI. Consistent with these observations, QKI levels increase in CM and decrease in DE, compared to undifferentiated hESCs, and disruption of QKI expression reveals that it is required for a substantial fraction of CM-specific splicing changes. One such target is exon 7 of *<u>B</u>ridging <u>IN</u>tegrator 1/Amphiphysin 2* (*BIN1*) transcripts. Through an unusual mechanism, inhibition of splicing of this exon by QKI is coupled to the promotion of adjacent intron removal, transcript maturation, and subsequent accumulation of BIN1 protein. Moreover, knockout of *QKI* prevents the transition from CM to cardiomyocytes, which likely is mediated in part by a requirement of *QKI* for expression of *BIN1*. Collectively, these findings uncover a key role for alternative splicing in defining early lineage-specific gene expression patterns, and implicate QKI as a cardiac mesoderm-enriched RBP that is required for the control of lineage-specific splicing and differentiation of cardiac myocytes.

## MATERIALS AND METHODS

### Cell culture and differentiation

All cells were cultured at 37°C. H9-hrGFP_NLS_ (a generous gift from Seigo Hatada), *NKX2.5*➔EGFP HES3 (a generous gift from David Elliot, Andrew Elefanty, and Ed Stanley (Murdoch Children’s Research Institute), and *NKX2.5*➔EGFP QKI KO human embryonic stem cells were cultured on hESC-qualified Matrigel (Thermo Fisher/Corning) and supplemented with mTeSR1 (Stemcell Technologies) supplemented with 1% penicillin/streptomycin (Thermo Fisher/Gibco). They were routinely passaged using ReLeSR or Gentle Cell Dissociation Reagent (both from Stemcell Technologies) following the manufacturer’s instructions and at times (such as recovery after cryopreservation) with the addition of Y27632 ROCK inhibitor (Thermo Fisher) according to the manufacturer’s instructions. Each of the hESC lines were derived from a female, were confirmed to have a normal karyotype during the course of the study, and routinely tested negative for mycoplasma contamination. hESC transfections were performed using Lipofectamine Stem Reagent (Thermo Fisher) according to the manufacturer’s instructions.

Differentiation of hESCs was performed essentially as previously described (7–10): cells were passaged at approximately a 1:5 to 1:20 ratio (depending on starting cell density and duration of differentiation experiment) and allowed to attached and equilibrate for 24-48 hours in mTeSR1 (Stemcell Technologies) supplemented with Y27632 ROCK inhibitor (Thermo Fisher). Then CDM2 basal media (50% IMDM (Thermo Fisher/Gibco), 50% F12 (Thermo Fisher/Gibco), 1 mg/ml polyvinyl alcohol (Sigma), 1% chemically defined lipid concentrate (Thermo Fisher/Gibco), 450 µm monothioglycerol (Sigma), 0.7 µg/ml insulin, 15 µg/ml transferrin) supplemented with 1% penicillin/streptomycin (Thermo Fisher/Gibco) was added for ECT differentiation. Media was changed daily and performed for three days. On day three, CDM2 was supplemented with 10% knockout serum replacement (Thermo Fisher/Gibco) and cultured an additional four days with daily media changes.

For endoderm differentiation cells were plated as above then first differentiated to anterior primitive streak for 24h in CDM2 basal media supplemented with 100 ng/ml Activin A (R&D Systems or Stemcell Technologies), 2 µm CHIR99021 (Tocris), and 50 nM PI-103 (Tocris), For the following 48h, cells were cultlured in CDM2 basal media supplemented with 100 ng/ml Activin A (R&D Systems or Stemcell Technologies) and 250 nM LDN-193189 (Tocris) with one media change after the initial 24h incubation.

For cardiac lineage differentiation cells were plated as above then first differentiated to mid primitive streak (MPS) for 24h in CDM2 basal media supplemented with 30 ng/ml Activin A (R&D Systems or Stemcell Technologies), 40 ng/ml BMP4 (R&D Systems or Stemcell Technologies), 6 µM CHIR99021 (Tocris), 20 ng/ml FGF2 (Thermo Fisher), and 50 nM PI-103 (Tocris). The following 24h were used to differentiated MPS to lateral mesoderm in CDM2 basal media supplemented with 30 ng/ml BMP4 (R&D Systems or Stemcell Technologies), 1 µM A-83-01 (Tocris), and 1 µM C59 (Thermo Fisher). For the next 24-48h lateral mesoderm was incubated in CM media (media changed every 24h), which consisted of CDM2 basal media supplemented with 30 ng/ml BMP4 (R&D Systems or Stemcell Technologies), 1 µM A-83-01 (Tocris), 20 ng/ml FGF2 (Thermo Fisher), and 1 µM C59 (Thermo Fisher). For cardiomyocyte differentiation, day four CM was supplemented with CDM2 containing 30 ng/ml BMP4 (R&D Systems or Stemcell Technologies), 1 µM XAV939 (Thermo Fisher), and 200 µg/ml 2-phospho-ascorbic acid (Sigma) for an additional 48-96h with daily media change.

Human embryonic kidney 293T/17 (HEK293T/17; ATCC) cells were obtained from the Duke University Cell Culture Facility. HuH7 WT control and QKI-KO#3 (39) cells were a generous gift from Kuo-Chieh Liao (Duke-NUS). Each of these was cultured in DMEM (Thermo Fisher/Lonza) supplemented with 10% FBS (Thermo Fisher/HyClone) and 1% penicillin/streptomycin (Thermo Fisher/Gibco) and routinely passaged with 0.25% Trypsin + 1 mM EDTA (Thermo Fisher/Gibco). HuH7 cells (WT and QKI KO#3) were transfected with Lipofectamine 2000 (Thermo Fisher) according to the manufacturer’s instructions. HEK293T/17 cells were transfected with polyethyenimine, linear MW 25 kDa (Polysciences, Inc.) as described (Addgene.org/protocols/transfection/).

### Plasmids and tranfections/transductions

*NKX2.5*➔GFP hESCs were co-transfected with pX458-sgQKI as described (39) or control empty vector along with pCEP-puro which contains a puromycin-resistance gene to allow for antibiotic selection and elimination of untransfected cells. One day later 1 μg/ml puromycin was added and cells were selected for 3 days, and then expanded in the absence of puromycin. Upon verification of reduced QKI protein in expanded cells, then directly sequencing PCR products amplified with PrimeSTAR Premix HS (Clontech) from total population-level genomic DNA (extracted using the DNeasy Blood and Tissue kit (Qiagen) by following the manufacturer’s directions) and analyzing with TIDE (40), limiting dilutions of 0.5 cells per well were performed and delivered to 96-well plates. These were monitored visually to confirm only single cells were present in each well, then expanded. Potential clonal KO lines were initially screened by western blot, then genomic DNA (extracted and amplified as above) was cloned into pCR4Blunt-TOPO (Thermo Fisher) then six different colonies derived from independent hESC clones were sequenced to confirm two copy disruption of the reading frame.

To construct inducible CRISPR-on *NKX2.5*➔EGFP cells, we co-transfected AAVS1-Neo-M2rtTA and AAVS1-idCas9-vpr with Lipofectamine Stem. One day later, we performed dual selection with 1 μg/ml puromycin plus 200 μg/ml G418 and maintained these conditions for 10 days. We confirmed doxycycline-dependent of Cas9:VPR by western blotting. For *QKI* induction, we cloned QKI sgRNA1 into pLenti SpBsmBI sgRNA Hygro and transfected this or the empty plasmid into HEK293T/17 cells along with packaging plasmid (pCMVR8.74) and VSV-G envelope plasmid (pMD2.G) to generate viral supernatant as described (addgene.org/protocols/lentivirus-production/) then concentrated with Amicon Ultra 100K centrifugal filter units (Millipore/Sigma).

Custom shRNA sequences (see Table S8) were generated and added to pLKO.1-hPGK-Neo (Millipore/Sigma) plasmid then viral supernatants were generated and collected as described above. Undifferentiated H9 hESCs were infected with equal volumes of lentivirus, then selected with 200 μg/ml G418 beginning 24h later and continuing for 48-72h, at which time the cells were harvested. Oligonucleotide sequences are listed in Table S8.

The pDUP51 splicing reporter plasmid has been previously described (41). To generate BIN1 exon 7 splicing reporters, exon 7 along with 126 bp of upstream and 184 bp of downstream flanking intronic sequence was PCR-amplified from human genomic DNA (extracted from WT *NKX2.5*➔EGFP cells) with PrimeSTAR Premix HS (Clontech) and ligated into pCR4Blunt-TOPO (Thermo Fisher). The WT DNA was sequenced to confirm the correct sequence, then the QuickChange^TM^ method was used to mutate putative exonic or intronic QKI motifs. These were again verified by sequencing then cloned into pDUP51 along with the WT sequence. The plasmids generated were pDUP-BIN1_ex7 (WT), pDUP-BIN1_ex7_EXdelMT (exon deletion mutant), pDUP-BIN1_ex7_EXsubMT (exonic substitution mutant), pDUP-BIN1_ex7_INTRdelMT (intronic deletion mutant), pDUP-BIN1_ex7_INTRsubMT (intronic substitution mutant), pDUP-BIN1_ex7_2xdelMT (exonic and intronic deletion mutant), and pDUP-BIN1_ex7_2xsubMT (exonic and intronic substitution mutant).

### Western blotting

Proteins were extracted using RSB100 buffer (100 mM Tris-HCl pH 7.4, 0.5% NP-40, 0.5% Triton X-100, 0.1% SDS, 100 mM NaCl, and 2.5 mM MgCl_2_) plus cOmplete EDTA-Free Protease Inhibitor Cocktail (Sigma), by direct addition of this buffer to wells/plates containing cells and scraped with cell scrapers and collect by pipet. Lysates were then vortexed intermittently while kept on ice for approximately 30 minutes, then centrifuged at 14,000 RPM at 4C for 15 minutes. Clarified lysate was transferred to a clean tube and either used immediately or store at −80C. Protein quantification was performed using the Bradford protein assay (BioRad). Equal protein amounts were loaded and separated on 10 or 12% SDS-PAGE then transferred to 0.4 μm nitrocellulose. Blocking was performed with 5% non-fat milk (NFM) in tris-buffered saline (TBS) for 1 hour at room temperature (RT). Primary antibody incubations (including anti-panQKI, anti-QKI5, anti-QKI6, anti-QKI7 (all four from Antibodies Incorporated), anti-QKI (Bethyl), anti-alpha/beta-Tubulin, anti-NKX2.5 (both from Cell Signaling Technologies), anti-SOX17 (R&D Systems), anti-OCT4, anti-cTNT, anti-BIN1 (all three from abcam), and anti-GAPDH (from Sigma) were performed overnight at 4C in NFM in TBS containing 0.01% tween-20 (TBST), then membranes were washed at least 3 times with TBST at RT. Secondary infrared-conjugated antibodies from Li-Cor were added to NFM in TBST at a dilution of 1:20,000 or 1:15,000, and incubated for 1 hour at RT protected from light. Membranes were then washed 3 times for 5 minutes and then 1-2 times for 15 minutes each before visualization on Odyssey CLx imager (Li-Cor).

#### Immunofluorescence

Cells were fixed with 4% paraformalydehyde diluted in phosphate-buffered saline (PBS) for 15-30 minutes then washed with PBS 2-3 times and stored in PBS until stained. Cells were blocked/permeabilized with 1% BSA and 0.3% Triton X100 in PBS for 1 hour at RT, and were then incubated overnight at 4°C in primary antibody (these included anti-panQKI (Antibodies Incorporated), anti-QKI (Bethyl), anti-NKX2.5, anti-SOX2 (both from Cell Signaling Technologies), anti-OCT4, anti-cTNT, anti-BIN1 (all three from abcam), anti-SSEA3, anti-SOX17 (both from R&D Systems), and anti-GAPDH (Sigma)) diluted in the above mentioned buffer. The following day, they were washed 3 times with PBS, then incubated in species-appropriate secondary antibody AlexaFluor 568 or AlexaFluor 647 (Thermo Scientific) at 1:1000 dilution in the above blocking buffer with the addition of 1 µg/ml DAPI) for 1 hour at RT, and then washed 3 times with PBS. Cells were then imaged immediately or stored for later analysis. Imaging was performed with Opera Phenix High Content Screening microscope (Perkin Elmer). Fluorescence intensity values were calculated for each channel on a per-cell and per-well basis using Harmony software (PerkinElmber), and recorded then subsequently analyzed in GraphPad Prism. Low magnification live cell imaging for NKX2.5➔EGFP shown in Fig S5F were performed on an IX71 inverted fluorescence microscope using the FITC filter (Olympus). Image analyses were done using Fiji (v2.0.0-rc-69/1.52p).

### RNA extraction, RT-qPCR, and RT-PCR

RNA was extracted from cells by directly adding TRIzol (Thermo Fisher) to wells containing cells, or using the ReliaPrep RNA MiniPrep system (Promega), each according to the manufacturer’s instructions. In the case of TRIzol-based extractions, nucleic acid was treated with Turbo DNase (Thermo Fisher) then RNA re-extracted with phenol/chloroform/isoamyl alcohol and chloroform extraction. Reverse transcription was performed with Super Script III Reverse Transcriptase (Thermo Fisher) using a combination of anchored oligo dT and random hexamers. Depending on the input material amount, cDNA was then diluted within a range of 1:10 to 1:50 then used for qPCR analysis with PowerUp SYBR Green Master Mix (Thermo Fisher) on a StepOnePlus Real-Time PCR machine (Thermo Fisher). For standard PCR, cDNA was added and amplified using Taq 2X Master Mix (NEB) with cycle number determined empirically to ensure no overamplification. PCR products were analyzed by either agarose gel or DNA1000 BioAnalyzer chip (Agilent) run on the 2100 BioAnalyzer (Agilent).

### RNA sequencing and analysis

RNA was extracted with TRIzol (see above; Thermo Fisher) and rRNA was depleted using the RiboCop rRNA Depletion Kit (Lexogen). This was used as input to generate paired-end libraries using the TruSeq Stranded Library Kit (Illumina) which were then pooled and sequenced on two High Output NextSeq 550 flow cells (Illumina) with a 75 base paired-end protocol. Reads were mapped using STAR version 2.6.1c onto the human GRCh38 reference build, with the parameters recommended by the ENCODE consortium. DESeq2, version 1.20.0, was used to determine changes in gene expression following the authors’ vignette. For splicing analyses, we used VAST-tools (version 2.0.2, database has.16.02.18) and rMATS (version 4.0.2, GENCODE v30 annotation) using the default parameters for each. Raw output was filtered based on quality scores, requiring at least 10 total reads per event in at least one replicate for a condition. Differential splicing was assessed with the VAST-tools diff module. Reads mapped onto the UCSC Genome Browser (hg38) were used to make coverage tracks and screen shots of these were used to visualize read levels for transcript abundance and alternative splicing. The RNAseq data was deposited at Gene Expression Omnibus (GSE162649).

### Computational analysis of CLIP and RNA-seq datasets

We collected functional genomic datasets from a variety of sources assaying RNA binding proteins, including ENCODE (42), CLIPdb (43), and starBase (44). All datasets were indexed by their genomic coordinates, which were used to compare to the genomic coordinates of alternative exons and 250bp of upstream or downstream introns using the RELI tool (45). In brief, RELI estimates the significance of the observed number of intersections between an input set of regions (e.g., alternative exon sites) and each member of a library of functional genomics datasets (e.g., CLIP-seq peaks). Significance is estimated by comparing to a null model formed by randomly sampling non-significant sites in each corresponding group used in this study (e.g., exons that are expressed (> 50 base mean of average normalized read counts) but do not significantly change in CM cell compared to DE cell dataset (P ≥ 0.5), and their flanking 250 bp of upstream and downstream intron sequence). The distribution of the expected intersection values from the null model resembles, and therefore is modeled as, a normal distribution, with the model parameters estimated from a series of 2,000 sampling procedures. To measure the frequency of RBP binding to alternative CE events, we merged the coordinates of each region of interest (250 bp upstream intron, cassette exon, and 250 bp of downstream intron) and performed RELI analysis as above. The “ratio” value indicates the binding frequency observed for a given RBP CLIP dataset. See below for statistical analysis used for RELI and RNA-seq.

### Flow cytometry and fluorescence-activated cell sorting (FACS)

Cells were prepared for FACS by removal from plates/wells with Accumax (Innovative Cell Technologies Inc.) according to the manufacturer’s instructions. Staining for CXCR4 surface marker was performed using APC anti-human CD184 (CXCR4; BioLegend) or with APC Mouse IgG_2a_, k isotype control (BioLegend) following the instructions from the manufacturer. The cells were finally passed through a 40 μm strainer then kept on ice before sorting with BD FACS Aria IIU (Beckton Dickinson) with assistance from core facility operator Mark Griffin. For analysis of EGFP levels in *NKX2.5*➔EGFP cells, cells were detached with 1x Trypsin/EDTA (0.25%; Thermo Fisher) according to the manufacturer’s instructions and either measured immediately or fixed with 2% paraformaldehyde dissolved in PBS if measured later on the Guava EasyCyte HT (Luminex) flow cytometer. For all flow/FACS applications, gating on relevant forward and side scatter parameters was performed to restrict any events not identified as single cells. FlowJo software was used for data analysis.

### Liquid chromatography with tandem mass spectrometry (LC-MS/MS)

Sample digestion: The samples were prepared similar to as described (46). Briefly, the agarose bead bound purified proteins were washed several times with 50mM TEAB pH 7.1, before being solubilized with 40uL of 5% SDS, 50mM TEAB, pH 7.55 and incubated at RT for 30 minutes. The supernatant containing proteins of interest was then transferred to a new tube and reduced by making the solution 10mM TCEP (Thermo, #77720) and incubated at 65^○^C for 10min. The sample is then cooled to RT and 3.75 uL of 1M iodoacetamide acid added and allowed to react for 20 minutes in the dark after which 0.5uL of 2M DTT is added to quench the reaction. 5 ul of 12% phosphoric acid is added to the 50uL protein solution. 350uL of binding buffer (90% Methanol, 100mM TEAB final; pH 7.1) is then added to the solution. The resulting solution is added to S-Trap spin column (protifi.com) and passed through the column using a bench top centrifuge (30s spin at 4,000g). The spin column is washed with 400uL of binding buffer and centrifuged. This is repeated three times. Trypsin is added to the protein mixture in a ratio of 1:25 in 50mM TEAB, pH=8, and incubated at 37^○^C for 4 hours. Peptides were eluted with 80uL of 50mM TEAB, followed by 80uL of 0.2% formic acid, and finally 80 uL of 50% acetonitrile, 0.2% formic acid. The combined peptide solution is then dried in a speed vac and resuspended in 2% acetonitrile, 0.1% formic acid, 97.9% water and placed in an autosampler vial.

NanoLC MS/MS Analysis: Peptide mixtures were analyzed by nanoflow liquid chromatography-tandem mass spectrometry (nanoLC-MS/MS) using a nano-LC chromatography system (UltiMate 3000 RSLCnano, Dionex), coupled on-line to a Thermo Orbitrap Fusion mass spectrometer (Thermo Fisher Scientific, San Jose, CA) through a nanospray ion source (Thermo Scientific). A trap and elute method is used. The trap column is a C18 PepMap100 (300um X 5mm, 5um particle size) from ThermoScientific. The analytical column is an Acclaim PepMap 100 (75um X 25 cm) from (Thermo Scientific). After equilibrating the column in 98% solvent A (0.1% formic acid in water) and 2% solvent B (0.1% formic acid in acetonitrile (ACN)), the samples (2 µL in solvent A) were injected onto the trap column and subsequently eluted (400 nL/min) by gradient elution onto the C18 column as follows: isocratic at 2% B, 0-5 min; 2% to 4% 5-6 min, 4% to 28% B, 6-120 min; 28% to 40% B, 120-124 min; 40% to 90% B, 124-126 min; isocratic at 90% B, 126-126.5 min; 90% to 5%, 126.5-127 min; 5%-90% 127-128min; 90%-5% 127-128min;and isocratic at 2% B, till 150 min.

All LC-MS/MS data were acquired using XCalibur, version 2.1.0 (Thermo Fisher Scientific) in positive ion mode using a top speed data-independent acquisition (DIA) method with a 3 sec cycle time. The survey scans (*m/z* 400-1000) were acquired in the Orbitrap at 60,000 resolution (at *m/z* = 400) in profile mode, with a maximum injection time of 50 msec and an AGC target of 4,000,000 ions. The S-lens RF level is set to 60. DIA windows were staggered at 12Da Isolation as described (47). HCD MS/MS acquisition is performed in profile mode using 15,000 resolution in the orbitrap, with the following settings: collision energy = 30%; maximum injection time 22 msec; AGC target 500,000 ions and scan range of 145-1450.

Database Searching: Each sample had three bioreps and the DIA data were analyzed using Scaffold DIA (3.0.1). Raw data files were converted to mzML format using ProteoWizard (3.0.21054) (48). Analytic samples were aligned based on retention times and individually searched against pan_human_library.dlib with a peptide mass tolerance of 10.0 ppm and a fragment mass tolerance of 10.0 ppm. The digestion enzyme was trypsin with a maximum of 1 missed cleavage site(s) allowed. Only peptides with charges of +2 or +3 were considered. Peptides identified in each sample were filtered by Percolator (3.01.nightly-13-655e4c7-dirty) (49–51) to achieve a maximum FDR of 0.01. Individual search results were combined and peptide identifications were assigned posterior error probabilities and filtered to an FDR threshold of 0.01 by Percolator (3.01.nightly-13-655e4c7-dirty).

Peptide quantification was performed by Encyclopedia (1.2.2). For each peptide, the 5 highest quality fragment ions were selected for quantitation. Only peptides exclusive to each protein or cluster were used for quantification. Proteins that contained similar peptides and could not be differentiated based on MS/MS analysis were grouped to satisfy the principles of parsimony. Proteins with a minimum of 2 identified peptides were thresholded to achieve a protein FDR threshold of 1.0%.

### Real-time bioenergetic analysis in intact cells

The XF24 Extracellular Flux Analyzer (Agilent, Santa Clara, USA) was used to measure bioenergetic function in intact, day 8 or day 15 cardiac-differentiated WT or QKI-KO (clone 4-2) HES3 NKX2.5➔EGFP as described previously (52, 53). In preliminary studies, we determined that 1.5 x 10^4^ cells was the optimal number of cells/well to allow detection of changes in oxygen consumption rate (OCR) for subsequent experiments. The XF24 Extracellular Flux Analyzer records the changes in oxygen levels in real-time by utilizing specific fluorescent dyes. Prior to the bioenergetic measurements, the culture medium was changed to unbuffered DMEM without serum. Next, we measured indices of mitochondrial function. Oligomycin, FCCP (carbonyl cyanide 4-(trifluoromethoxy) phenylhydrazone), and antimycin A/rotenone (AA + Rot) were injected sequentially through ports of the Seahorse Flux Pak cartridges to reach 2 µM, 1 µM, 2 µg/ml and 2 µM, respectively. Key bioenergetic parameters such as basal respiration (resting cell respiration), ATP production (calculated from the drop in OCR, in response to the ATP-synthase inhibitor, oligomycin), proton leak (migrated protons to the matrix without producing ATP), maximal respiratory capacity (maximal oxygen consumption achievable by using the uncoupling agent FCCP), spare respiratory capacity (accessible mitochondrial reserve capacity under high bioenergetic demands), and non-mitochondrial O_2_ consumption (a small amount of residual OCR related to O_2_ consumption due to the formation of reactive oxygen species or function of extramitochondrial enzymes) were measured. All measurements were normalized to protein content, determined in each individual well.

#### Reagents

MouseIgG_2b_-anti-panQKI (Clone N147/6; cat# AB_2173149), mouseIgG_1_-anti-QKI5 (Clone N195A/16; Cat# AB_10674117), mouseIgG_1_-anti-QKI6 (Clone N182/17; Cat# AB_10673512), mouseIgG_1_-anti-QKI7 (Clone N183/15; Cat# AB_2173148) are from Antibodies Incorporated/Millipore (Burlington, MA USA), Rabbit-anti-QKI (Cat# A300-183A) is from Bethyl/Fortis (Waltham, MA USA), Rabbit-anti-alpha/beta-Tubulin (Cat# 2148), Rabbit-anti-NKX2.5 (Clone E1Y8H; cat# 8792S), mouseIgG_1_-anti-SOX2 (L1D682; cat# 4900S) are from Cell Signaling Technology (Danvers, MA USA), mouseIgG_2b_-anti-OCT4 (Cat# ab184665), Rabbit anti-BIN1 (Cat# ab182562), and rabbit-anti-cTNT (Cat# ab91605) are from abcam (Cambridge, UK) and Goat-anti-SOX17 (Cat# AF1924), ratIgM-anti-SSEA3 (Clone MC-38; cat# MAB1434) are from R&D Systems (Minneapolis, MN USA), mouseIgM-anti-GAPDH (Clone GAPDH-71.1; cat# G8795) is from Sigma-Aldrich (St. Louis, MO USA), mouseIgG_2a_-anti-CD184, APC conjugate (Clone 12G5) and mouseIgG2a isotype, APC conjugate (Clone MOPC-173) are from BioLegend (San Diego, CA USA), Guide-it™ Cas9 Polyclonal Antibody (Cat#632607) and PrimeSTAR Premix HS (Cat# R010A) are from Takara Bio Inc (Kusatsu, Shiga, Japan), Goat anti-Mouse IgG2b Cross-Adsorbed Secondary Antibody, AlexaFluor 568 (Cat #A-21144), Goat anti-Mouse IgG1 Cross-Adsorbed Secondary Antibody, AlexaFluor 647 (Cat# A-21240), Goat anti-Rabbit IgG (H+L) Cross-Adsorbed Secondary Antibody, AlexaFluor 568 (Cat# A-11011), Donkey anti-Goat IgG (H+L) Cross-Adsorbed Secondary Antibody, AlexaFluor 647 (Cat# A-11057), Donkey anti-mouse IgG (H+L) Cross-Adsorbed Secondary Antibody, AlexaFluor 568 (Cat# A-10037), and Goat anti-Rat IgG (H+L) Cross-Adsorbed Secondary Antibody, AlexaFluor 647 (Cat#A-21247), PowerUP SYBR Green Master Mix (Cat# A25742), Lipofectamine 2000 Transfection Reagent (Cat# 1166027), Lipofectamine Stem Reagent (Cat# STEM00015), Super Script III Reverse Transcriptase (Cat# 18080085), pCR4Blunt-TOPO (Cat# K2875J10), TRIzol Reagent (Cat# 15596026), and Turbo DNase (Cat# AM2238) are from Thermo Fisher (Waltham, MA USA). RiboCop rRNA Depletion Kit (Cat# 037.24) is from Lexogen (Wien, Austria). Tru Seq Kit (Cat# RS-122-2101) is from Illumina (San Diego, CA USA). BioAnalyzer DNA1000 Kit (Cat# 5067-1505) is from Agilent (Santa Clara, CA USA). Amicon Ultra 100K centrifugal filter units (Cat# UFC910008) are from Millipore Sigma (Burlington, MA USA). ReliaPrep RNA Cell Miniprep System (Cat# Z6012) is from Promega (Madison, WI USA). Taq 2x Master Mix (Cat# M0270L) is from New England Biolabs. The DNeasy Blood and Tissue Kit (Cat# 69504) is from Qiagen (Hilden, Germany). The Bradford Protein Assay Kit (Cat# 5000201) is from BioRad (Hercules, CA USA). Activin A (Cat# 33AC050 or 78001.1), BMP4 (Cat# 314-BP-050 or 78211), FGF2 (Cat# PHG0023), Chemically defined lipid concentrate (Cat# 1905-031), CHIR99021 (Cat# 44-231-0), Knockout Serum Replacement (Cat# A3181502), WNT-C59, PORCN INHIBITOR II (Cat# NC1022261), XAV939 (Cat# 50-152-9985), Y27632-ROCK Inhibitor (Cat# NC0407157), IMDM (Cat# 31-980-030), F12 (Cat# 31-765-035), DMEM (Cat# BW12604F12), FBS (Cat# SH3007003HI), 0.25% Trypsin + 1 mM EDTA (1x Trypsin/EDTA) (Cat#25200056), Penicillin/Streptomycin (Cat# 15140122), and Corning Matrigel hESC-Qualified Matrix are from Thermo Fisher (Waltham, MA USA). Polyvinyl alcohol (Cat# P8136-1KG), Monothioglycerol (Cat# M6145), 2-phospho-ascorbic acid (Cat# 49752-10G), Insulin, human (Cat# I2643-50MG), Holo-transferrin, human (Cat# T0665-100MG), and cOmplete, EDTA-free protease inhibitor cocktail (Cat# 11873580001) are from Sigma Aldrich (St. Louis, MA USA). PI-103 (Cat# 2930), LDN-193189 (Cat# 6053), A-83-01 (Cat# 2939), are from Tocris (Bristol, UK). Accumax (Cat# AM105) is from Innovative Cell Technologies, Inc. (San Diego, CA USA). mTeSR1 (Cat# 85857), ReLeSR (Cat# 05873), and Gentle Cell Dissociation Reagent (Cat# 07174) are from Stemcell Technologies (Vancouver, Canada). Polyethyenimine, linear MW 25,000 Da (Cat# 23966-1) is from Polysciences, Inc. (Warrington, PA USA).

### Biological Resources

Cells and cell lines used: H9-hrGFP_NLS_ human embryonic stem cells are from Seigo Hatada (54), *NKX2.5*➔eGFP HES3 human embryonic stem cells are from Andrew Elefanty and Edward Stanley (55), the QKI knockout *NKX2.5*➔eGFP HES3 and the dCas9:VPR knock-in *NKX2.5*➔eGFP HES3 human embryonic stem cells were generated in this study. HuH7 QKI-KO#3 and wild type control cells are from Kuo-Chieh Liao (39). HEK293T/17 are from ATCC (CRL-11268).

Plasmids and other recombinant DNA: pX458-sgQKI was a gift from Kuo-Chieh Liao (39), pCEP-Puro was a gift from Namrita Dhillon, pMD2.G (Addgene plasmid #12259) and pCMVR8.74 (Addgene plasmid #22036) were from Didier Trono, pDUP51 was a gift from Ryszard Kole (41), pDUP-BIN1_ex7, pDUP-BIN1_ex7_EXdelMT, pDUP-BIN1_ex7_EXsubMT, pDUP-BIN1_ex7_INTRdelMT, pDUP-BIN1_ex7_INTRsubMT, pDUP-BIN1_ex7_2xdelMT, pDUP-BIN1_ex7_2xsubMT were generated in this study, pLK0.1-hPGK-Neo is from Sigma (https://www.sigmaaldrich.com/deepweb/assets/sigmaaldrich/marketing/global/documents/387/326/plko1-hpgk-neo.pdf), AAVS1-idCas9-vpr (Addgene plasmid 89985) is from Jie Na, pLenti SpBsmBI sgRNA Hygro (Addgene plasmid 62205) is from Rene Maehr, AAVS1-Neo-M2rtTA (Addgene plasmid 60843) is from Rudolph Jaenisch.

### Statistical analyses

Error bars used in any figures represent standard deviation (SD), and asterisks indicate the statistical significance as follows: \**P* < 0.05, \*\**P* < 0.01, \*\*\**P* < 0.001, \*\*\*\**P* < 0.0001. These were calculated using Prism8 or Prism9 (GraphPad). All Student’s t-tests were calculated using the two-sided test.

For quantification of protein abundances by western blot, after scanning infrared signals on the Odyssey CLx (Li-Cor), the region of interest for each band was selected and the value recorded using Image Studio software (Li-Cor). The values measured were normalized to a “loading control” value (either Tubulin or GAPDH), at which time the Student’s t-test was used to determine if experimental treatments were statistically significant between groups. In some cases, these normalized protein abundances were converted to values relative to control by dividing each measured experimental replicate value by the mean control value to determine the ratio of experimental to control; mean control values were then displayed at a value of 1.0 with experimental mean values shown relative to control, each +/-SD.

RT-qPCR Data were analyzed using the delta-delta CT method and reported as relative to *HMBS* “housekeeping gene”; we also confirmed similar results using values obtained from *EEF1A1* and *GAPDH* as “housekeeping genes”. Statistical significance was measured using the Student’s t-test with dCT values from biological replicates as input. In most cases data were displayed ddCT (2^^-dCT^) values as these convey dynamic changes in various cellular states, e.g. during differentiation between lineages or progeny cell types. In some cases data were displayed as relative to control, as described above.

RT-PCR/BioAnalyzer quantification was determined by calculating the ratio of PCR products’ concentration (nM) and reporting mean percent included or product represented of the band(s) of interest +/- SD.

Quantification of immunofluorescence data was performed with Harmony software (PerkinElmer). No correction was performed for very low signal as some is detectable in any given channel due to background, and as such, this was included in reported values. Population-level quantification (signal per well) was measured for statistical significance by directly comparing WT to KO signal (experimental signal (e.g. QKI, BIN1, etc.) was normalized to either DAPI or GAPDH signal) output with the Student’s t-test. Quantification of fluorescence intensity signals on the per cell level was normalized as above (using DAPI or GAPDH) for each event, then these values were binned and a histogram of distribution was generated. Statistical significance was measured by the Mann–Whitney U test.

Splicing events calculated by VAST-tools were deemed significantly differential if they had a 95% likelihood of being differential (i.e., MV[abs(dPSI)_at_0.95] > 0) and the mean PSI/PIR difference was greater than 10. Changes between lineages were displayed in violin plots and statistical significance between distributions of positive/negative events were measured by Mann–Whitney U test. GO analysis was done separately for CE/MIC and IR event types with a background comprising all genes that contain a splicing event of the type being tested that survived filtering, requiring a minimum three-fold enrichment. If categories mutually overlapped by more than 70%, only the more significantly enriched category was kept.

For RELI analysis the significance of the observed number of intersections, e.g., a Z-score and the corresponding P-value, along with an enrichment score (observed intersection count divided by expected intersection count) were calculated, and are provided in Table S4. We enlisted a significance cutoff of adjusted P-value < 0.05, ratio > 0.05, and enrichment ≥ 2 to ensure any given RBP’s binding was well-represented in our splicing datasets.

Quantification of EGFP levels by flow cytometry were determined by analysis using FlowJo software and the histogram output was also generated by this program. Statistical significance between mean values were measured between independent biological replicates and calculated with the Student’s t-test.

### Data Availability

The RNAseq data was deposited at Gene Expression Omnibus (GSE162649) and can be accessed with reviewer token ulirkgwkdfqvrcv; proteomics datasets were deposited at MassIVE (MSV000088762), and reviewers can access using password UTMBsfagg and username MSV000088762.

### Programs, Software, Algorithms, and Websites

STAR v2.6.1c https://github.com/alexdobin/STAR, VAST-tools v2.0.2 https://github.com/vastgroup/vast-tools#combine-output-format, rMATS v4.0.2 https://github.com/Xinglab/rMATS-STAT, DESEQ2 v1.20.0 https://github.com/mikelove/DESeq2, Fiji v2.0.0-rc-69/1.52p https://imagej.net/Fiji, Prism8 v8.4.3 GraphPad, FlowJo (BD Biosciences) https://www.flowjo.com/solutions/flowjo, Harmony v4.9 (PerkinElmer), UCSC Genome/Table Browser Hg38 genome.ucsc.edu, Tracking Indels by Decomposition (TIDE) tide.nki.nl (40), Image Studio Software V5.x (Li-Cor), RELI Github.com/weirauchlab/RELI#, and rMAPS2 http://rmaps.cecsresearch.org/ (56).

## RESULTS

### Identification of developmentally-regulated and lineage-specific alternative splicing

To identify alternatively spliced events in human pluripotent stem cells and their germ layer-committed progenitor cells, we differentiated H9 hESCs (undifferentiated, UD) in parallel to DE (7), CM; (8), and ECT (Fig 1A; (9)), and performed bulk RNA-sequencing (RNA-seq) to analyze their transcriptomes. We first measured known pluripotency- or lineage-specific biomarkers at the level of RNA (*POU5F1*/OCT4 for UD, *SOX17* for DE, *HAND1* for CM, and *PAX6* for ECT; see Fig 1B) and protein (CXCR4 for DE, see Fig S1 A; SOX2 and SSEA for UD, SOX17 for DE, NKX2.5 for CM, and SOX2 for ECT, see Fig S1 B), which indicate specific and efficient differentiation, consistent with previous reports (7, 8). Additionally, principal component analysis of transcript level changes (57) shows a close clustering of the three sample replicates from each lineage, indicating a high level of reproducibility (Fig S1 C). As noted previously (13) most protein-coding transcript abundance changes relative to UD (*P* ≤ 0.01 by Benjamini-Hochberg) were shared between germ layer-committed progenitor cells (Fig S1 D; Table S1).

**FIGURE 1:**
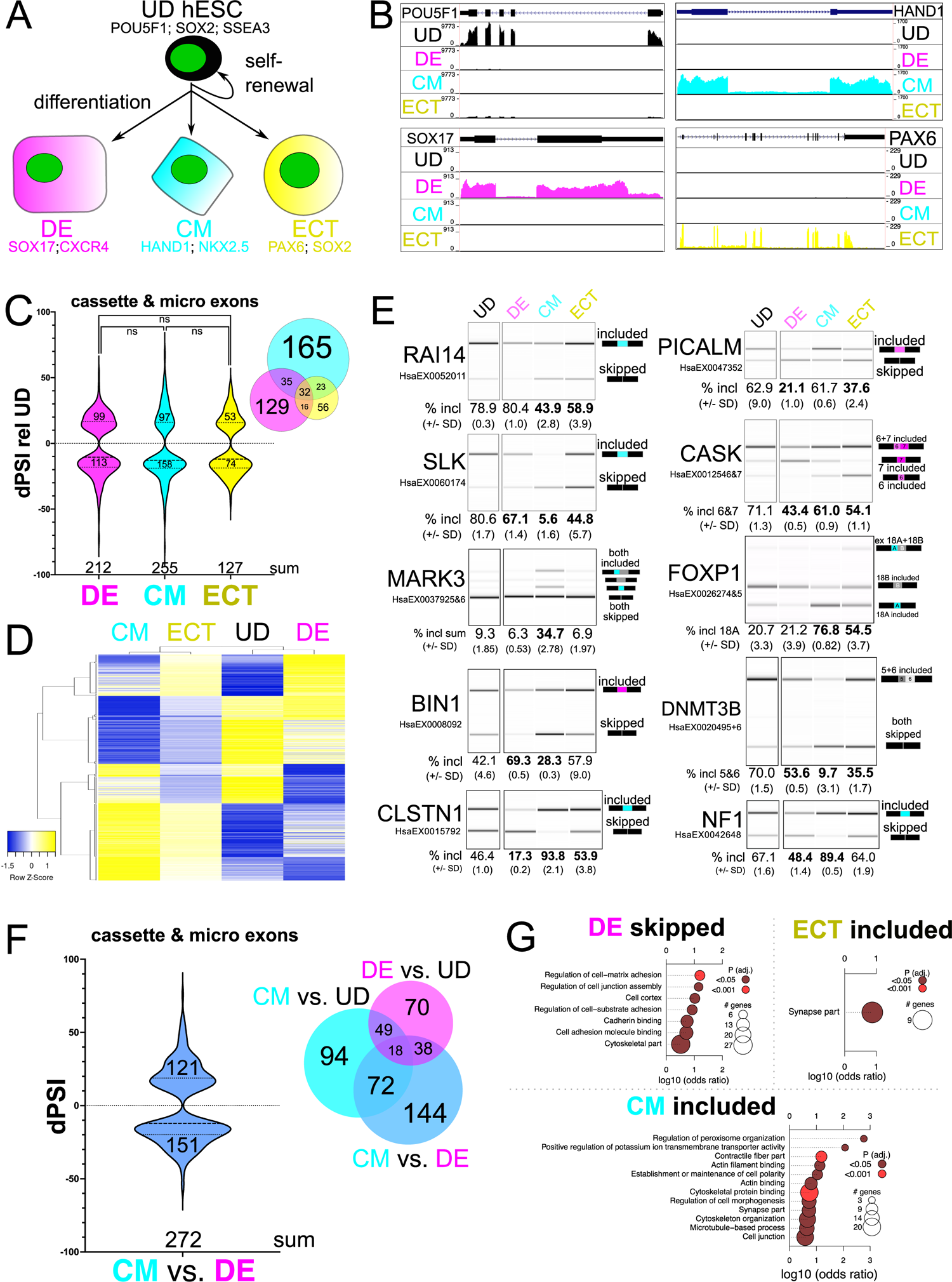
Identification of lineage-specific alternative splicing. **A.** Experimental overview/model of cell lineages analyzed by RNA-seq: H9 undifferentiated (UD) hESCs can self-renew or were differentiated to definitive endoderm (DE), cardiac mesoderm (CM), or neuroectoderm (ECT) cells for further analysis. Lineage-specific biomarkers are shown for each progenitor cell type: *POUF51*, *SOX2* and *SSEA3* for UD hESCs, *SOX17* and *CXCR4* for DE cells, *HAND1* and *NKX2.5* for CM cells, and *PAX6* and *SOX2* for ECT cells. **B.** UCSC Genome Browser screen shots showing coverage tracks for reads from RNA-seq representing the cell lineages described above, measuring RNA abundance for *POU5F1*/OCT4, *SOX17*, *NKX2.5*, and *PAX6*. **C.** Change in percent spliced in (dPSI) of alternative cassette and micro exons for DE cells, CM cells, and ECT cells, significantly changing relative to UD hESCs shown in violin plot, and overlap of these shown in Venn diagram inset, obtained from VAST-tools analysis of RNA-seq data (ns indicates no significant change by Mann–Whitney U). **D.** Heatmap showing hierarchical clustering of CEs that change significantly in at least one lineage; PSI values from UD, DE, CM, and ECT were used to make the heatmap and are displayed as row Z-score values. **E.** Validation by RT-PCR and BioAnalyzer quantification of alternatively spliced exons (VAST-tools alternative exon designation shown below transcript name) using RNA extracted from UD hESCs, DE cells, CM cells, and ECT cells (mean percent included +/- standard deviation from the mean is shown below; values that are statistically significant by Student’s t-test (upper bound is P ≤ 0.05 by Student’s t-test) compared to UD are shown in bold). **F.** dPSI of alternatively spliced cassette and micro exons that significantly changed in CM cells relative to DE cells shown in violin plot, and the overlap of these with CM cells compared to UD hESCs and DE cells compared to UD hESCs is shown in the Venn diagram inset; obtained from VAST-tools analysis of RNA-seq data. **G.** Gene ontology (GO) enrichment of genes with significantly changing alternative cassette/micro exons in CM cells, DE cells, or ECT cells relative to UD hESCs. See also Fig S1.

We used VAST-tools (58, 59) to measure splicing changes in each lineage relative to UD hESCs and detected 588 events in ECT, 977 in DE, and 1,144 in CM (dPSI ≥ 10, see Methods); Fig S1 E and Table S2). The majority of these are alternative cassette exons or 3-27 nt microexons (CE/MIC), or retained introns (Fig S1 E). We observe a similar frequency of increased inclusion vs skipping among differentially spliced CE/MIC events in DE, CM, and ECT, compared to UD cells (Fig 1C). The majority of alternatively spliced exons detected in DE or CM cells are specific to either lineage (unlike for transcript abundance changes (Fig S1 D)), but more than half of the events detected in ECT cells overlap with the other lineages (Fig 1C) irrespective of the cutoffs used (i.e. lower or higher dPSI values (Table S2)). We observe a similar trend for intron retention (IR), however there are significantly more retained introns in CM cells and fewer in DE or ECT cells (*P* < 0.0001 by Mann–Whitney U); additionally the frequency of unique IR events in ECT cells is higher compared to CE/MIC events in ECT cells (Fig S1 F). Furthermore, hierarchical clustering of CE/MIC (Fig 1D) or IR (Fig S1 G) events revealed that CM and ECT splicing patterns are more similar to each other than to DE. These observations reveal that most alternatively spliced exons detected in DE and CM cells are spliced in a lineage-specific manner.

We validated exons strongly differentially spliced in either CM cells (*RAI14* exon 11, *SLK* exon 13, and *MARK3* exons 17 and 18) or DE cells (*PICALM* exon 17 and *CASK* exon 19) by RT-PCR (Fig 1E). Interestingly, alternative exons in *FOXP1* (exons 18A or 18B; (60)) and *DNMT3B* (exons 21 and 22; (61–63)), previously defined for their roles in controlling transitions between pluripotent and differentiated states, show relatively large inclusion level changes in CM cells, and to a lesser extent in ECT cells or DE cells relative to UD hESCs (Fig 1E). An additional class of exons shows opposite splicing patterns between DE and CM cells; this class includes *BIN1* exon 7, which, relative to UD cells is more included in DE cells, more skipped in CM cells, and relatively unaffected in ECT cells. Other examples such as *CLSTN1* exon 11 and *NF1* exon 23 show the opposite trend (Fig 1E). Validation of these specific events further corroborates that DE and CM splicing patterns are the most divergent groups compared here (Fig 1D). Direct comparison using VAST-tools of CM relative to DE revealed 1,382 total alternatively spliced events (Fig S1 E), including 144 additional unique alternative exons (Fig 1F, compare to Fig 1C) and 682 distinct retained introns (Fig S1 H) not detected when compared to UD hESCs. Many of these are due to a modest alteration in inclusion in DE relative to UD, and an additional but opposite modest change in inclusion in CM relative to UD (Table S2). Moreover, specific CE/MIC events detected by RNA-Seq were validated at a high rate by RT-PCR assays (dPSI correlation: DE R^2^ = 0.87, CM R^2^ = 0.94, ECT R^2^ = 0.74, DE/CM R^2^ = 0.98), supporting the robustness of the measurement (Fig S1 I). We also observed significant enrichment of functional terms associated with genes containing these differentially spliced exons that are consistent with the biology of each lineage: i.e., the term synapse part is enriched for ECT cells, various cell adhesion/junction-based terms are enriched in genes with more skipping in DE cells, and contractile fiber part and related terms are associated with genes displaying increased exon inclusion in CM cells, (Fig 1G). Finally, we used rMATS (64) to independently identify CEs in each of the above comparisons and intersected these with exons significantly alternatively spliced by VAST-tools analysis. This indicated a positive correlation between the two dataset outputs, and a similar pattern of overlap between lineages that we observed with VAST-tools CEs (Fig S1 J; Table S3), comparable to previous observations (65). Collectively, these data suggest that alternative splicing in germ layer-committed progenitor cells represents a lineage-specific regulatory program that is orthogonal to transcriptional regulation.

### Binding and motif enrichment analyses reveal a role for QKI in CM-specific alternative splicing

We hypothesized that lineage-specific alternative splicing is regulated by distinct RBPs, and tested this with multiple approaches. First, we re-purposed a transcription factor/chromatin data analysis method called <u>R</u>egulatory <u>E</u>lement <u>L</u>ocus Intersection or RELI (45) to measure RBPs binding in or around lineage-specific CE/MIC events (CM compared to DE) identified as significantly differentially spliced by both VAST-tools and rMATS outputs (Fig S1 H and Tables S2 and S3). This generated a list of 68 skipped and 62 included exons. In brief, RBP-RELI uses a simulation-based procedure to systematically gauge the significance of the intersection between a set of input regions (e.g., and in this case, exons alternatively-spliced between CM and DE cells, together with 250bp of flanking upstream and downstream intron sequence) and each member of a large library of functional genomics experiments, in this case, peaks from over 450 RBP crosslinking immunoprecipitation (CLIP)-seq experiments ((43,44,66). This measures the binding frequency (fraction of splicing events), enrichment (fold-change over background), and significance (p-value) of RBPs to the lineage-specific CEs and their flanking intronic regions, compared to a background set of expressed exons and introns whose splicing is unchanged. This approach limits experimental bias arising from focusing on a single RBP, and facilitates discovery-based analysis.

RBP-RELI analysis yielded a list of 55 total unique CLIP datasets representing 17 individual RBPs with significant binding in or around exons that are alternatively spliced in CM cells compared to DE cells (cutoff: P-adj < 0.05, ratio > 0.05, enrichment ≥ 2 (see Methods); Table S4). We plotted the resulting z-scores of the top 10 most significant (or fewer if observed) RBPs relative to their transcript binding region (upstream intron, exon, downstream intron) and whether higher inclusion or skipping was observed (Fig 2A). This analysis indicated that Quaking (QKI) is the most frequently observed RBP with CLIP peaks mapping in or around these lineage-specific CEs (14 occurrences out of 55 total significant RBP datasets; Fig 2A and Table S4). Moreover, QKI may directly bind up to 45% of transcripts alternatively spliced between CM cells and DE cells (measured by at least one CLIP peak mapping within the genomic coordinates encompassing the CE and its flanking intron sequences (Fig S2 A and Table S4)). The most significant binding was observed in the exon or upstream intron associated with skipping, or in the downstream intron when associated with inclusion, consistent with previously described positional influences of QKI on alternative splicing (Fig 2A; (29, 67)). We also used the output comparing alternatively spliced exons in CM cells to DE cells by rMATS to measure motif enrichment with rMAPS2 (56). Consistent with the RBP-RELI results, this showed significant (by Wilcoxon’s rank sum test) position-dependent enrichment of the QKI binding motif ACUAA that correlated with either exon inclusion or skipping (Fig 2B and Table S5) (68–70). Specific examples are: CM cell-activated exons in *NF1*, *CLSTN1*, and *MARK3* (Fig 1E), which show QKI binding in the downstream intron (Fig 2C and S2B); CM cell-repressed exons in *RAI14* and *SLK* (Fig 1E) show QKI binding in the upstream intron and/or exon (Figs 2C and S2B); the DE cell-repressed exon in *PICALM* (Fig 1E) shows QKI binding in the downstream intron (Fig S2 B). Additional observations suggest a role for TIA1 and TIAL1 in promoting inclusion by binding introns downstream of these lineage-specific CEs, and for RBFOX2 acting as a repressor of these exons (Fig 2A). This was also apparent from the rMAPS2 analysis; additionally, we observed enrichment of the CU-rich PTB family binding motif, which was particularly associated with exon skipping (Table S5). Collectively, these results highlight the use of RBP-RELI to effectively identify candidate splicing regulatory RBPs of alternative splicing programs and suggest a prominent role for QKI in the regulation of CM-specific alternative splicing.

**FIGURE 2:**
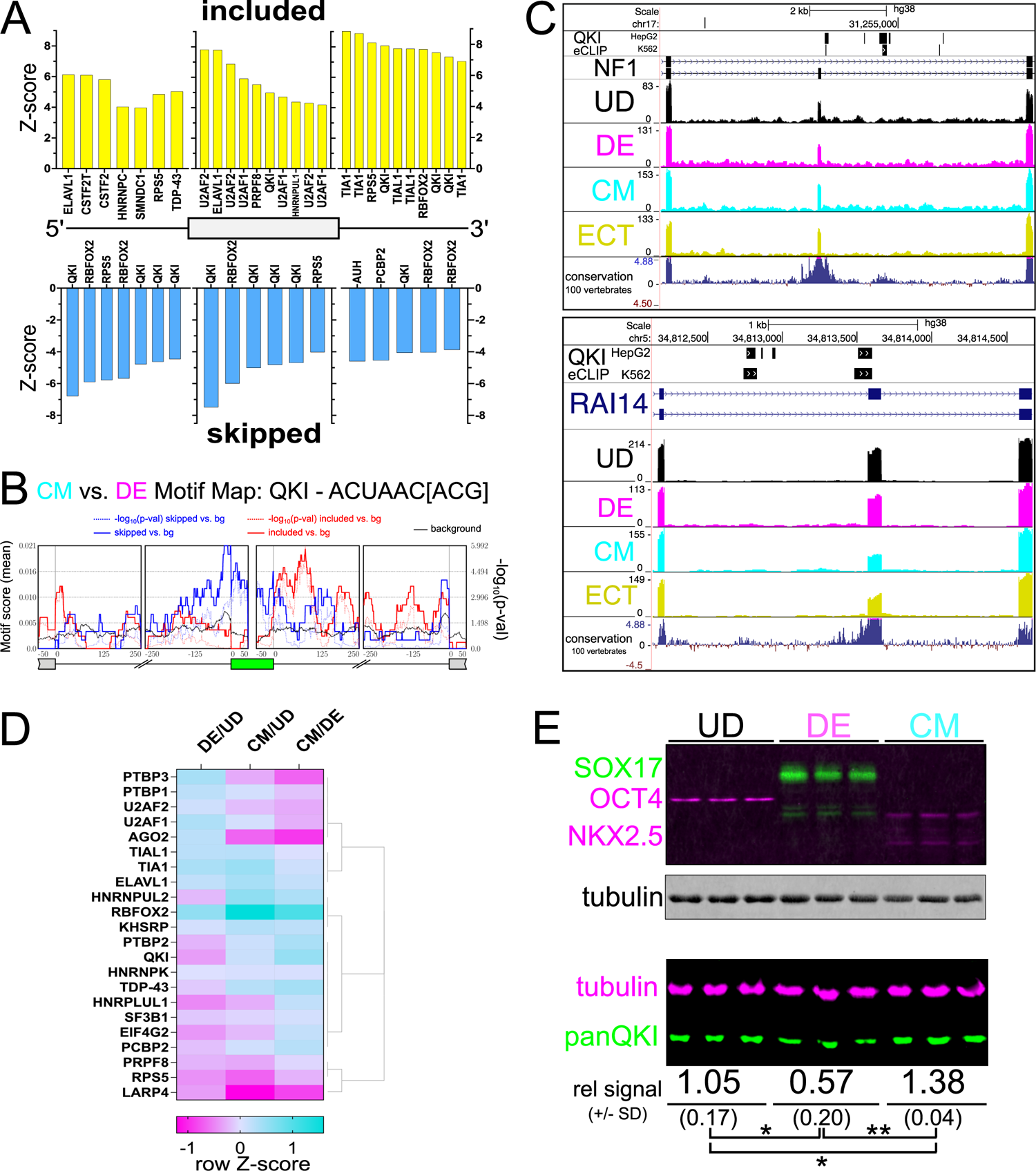
Significant binding and motif enrichment for QKI in CM-specific alternatively spliced exons, and QKI5 increases in abundance during CM differentiation. **A.** Enrichment (Z-score) of RBP eCLIP peaks mapped onto alternative exons either more included (above zero on y-axis in yellow) or more skipped (below zero on y-axis in blue) in CM cells vs. DE cells and mapped positionally by upstream intron (left) within alternative exon (middle) or in the downstream intron (right), calculated by RELI analysis of cassette exons identified as significantly differentially spliced in both VAST-tools and rMATS analyses of RNA-seq data. Each instance of a specific CLIP dataset that significantly correlated with CM cells vs. DE cells alternatively spliced exons is shown, therefore there may be more than one representation of a given RBP per region. **B.** RNA motif map (from rMAPS2) for QKI derived from rMATS analysis of alternatively spliced exons significantly changing when comparing CM cells relative to DE cells. **C.** UCSC Genome Browser screen shots showing QKI eCLIP peaks (HepG2 top, K562 below), Gencode Gene annotation for *NF1* (top) and *RAI14* (bottom), with RNA-seq coverage tracks for UD (black), DE (magenta), CM (cyan), and ECT (yellow), and conservation of 100 vertebrates track. See also Fig S2 A. **D.** Heatmap showing hierarchical clustering of protein abundance measurements by LC-MS/MS of annotated RBPs that were significantly represented in RELI (see Fig 2A) and/or rMAPS2 (see Table S5) analyses in DE cells or CM cells relative to UD hESCs, or CM cells relative to DE cells; relative abundance is shown as row Z-score. **E.** Western blot of protein extracted from triplicate cultures of UD hESCs, DE cells, and CM cells probed with antibodies for (top) OCT4 (magenta), SOX17 (green), NKX2.5 (magenta), and tubulin (gray), or (bottom) probed with antibodies for tubulin (magenta) and panQKI (green), showing mean infrared signal +/- standard deviation for panQKI relative to tubulin (rel IR signal +/- SD) below (\**P* ≤ 0.05, \*\**P* ≤ 0.01 by Student’s t-test).

### QKI5 increases in CM cells and decreases in DE cells

Given the above results, we hypothesized that QKI and possibly other splicing regulatory RBPs’ levels increase during CM- and decrease during DE-lineage commitment. We identified RBPs (71) whose abundance significantly change (base mean of average normalized read counts > 50; *P* ≤ 0.01 by Benjamini-Hochberg; log_2_ fold change > 0.4) (57) in our RNA-seq data (Table S1) and found that each germ layer representative cell type has a unique network of RBPs, with comparison of CM to DE cells representing the most divergent set (Fig S2 C). Likewise, comparing the proteomes of DE, CM, and UD cell extracts by liquid chromatography/tandem mass spectrometry (LC-MS/MS) indicated the most unique set of RBPs is detected when comparing CM to DE (Fig S2 D; Table S6). This provides further evidence suggesting that lineage-enriched RBPs direct lineage-specific splicing, and allowed for rapid identification of RBPs: 1) whose levels change in CM compared to DE, and 2) that show significant binding and/or motif enrichment in the CM cell compared to DE cell splicing datasets. We focused on this short list of candidates and found both QKI and PTBP2 increase in CM and decrease in DE cells at both the RNA (Table S1) and protein (Fig 2D) level. Of these, QKI exhibited significant binding (PTB family members failed to pass statistical cutoff; Fig 2A) and more binding (45% of significant CEs for QKI in HepG2 replicate 1 compared to 37% for PTBP1 in HepG2 replicate 2; Table S4 and Fig S2 A) by RELI analysis, and more significant and widespread motif enrichment (lowest P = 1.02^-6^ for QKI compared to 7.72^-6^ for PTBP1 by Wilcoxon’s rank sum test; Fig 2B and Table S5) to CM-specific alternatively spliced exons. Given these observations we chose to focus on QKI-regulated splicing during the exit from pluripotency and commitment to DE or CM. Western blot analysis of pluripotency (OCT4), DE (SOX17), or CM (NKX2.5) differentiation markers indicate specific and efficient differentiation of each cell type, and validated the changes observed for QKI protein by LC-MS/MS analysis (Fig 2E). Specifically, QKI significantly decreased in DE and increased in CM cell extracts, relative to UD hESCs (*P* < 0.05 by Student’s t-test; Fig 2E). Consistent with a greater magnitude of change in QKI splicing targets, comparison of its levels from CM cell lysates to DE cell lysates indicated an 81% increase in CM cells (**P < 0.01 by Student’s t-test; Fig 2E). We also measured the abundance of QKI5, QKI6, and QKI7 isoforms relative to total QKI protein as previously described (33), and found that the most abundant (>95%) is the nuclear isoform QKI5 (Fig S2 E). Additionally, we verified that QKI protein is predominantly nuclear, and co-expressed with lineage-specific biomarkers (SSEA3 for UD, SOX17 for DE, and NKX2.5 for CM; Fig S2 F). Collectively, these data support a prominent and direct role for QKI in the regulation of an alternative splicing program specifically linked to formation of the CM lineage.

### Cardiac mesoderm-specific exons are responsive to the levels of QKI

To determine whether QKI promotes CM-specific alternative splicing, we used CRISPR/Cas9 editing to generate hESC QKI knockout (QKI-KO) cells expressing EGFP under the control of the endogenous *NKX2.5* promoter, which is specifically activated in cardiac mesoderm/myocytes (55), and then measured the inclusion levels of several CM-specific alternatively spliced exons in this line. Disruption of the QKI ORF was performed using a guide RNA targeting exon one as previously described (39) and confirmed by analysis of PCR products derived from genomic DNA (Fig S3 A). We also selected and expanded single cells from which three clonal *NKX2.5*➔GFP knockout hESCs were assessed. Clones 1-12 and 4-2 have the same two lesions, while clone 4-5 shares one of the same lesions and an additional unique lesion (Fig S3 A). QKI protein is undetectable in all three clonal and the mixed population of cells (Fig 3A). Correspondingly, the inclusion of predicted QKI target exons changes.

**FIGURE 3:**
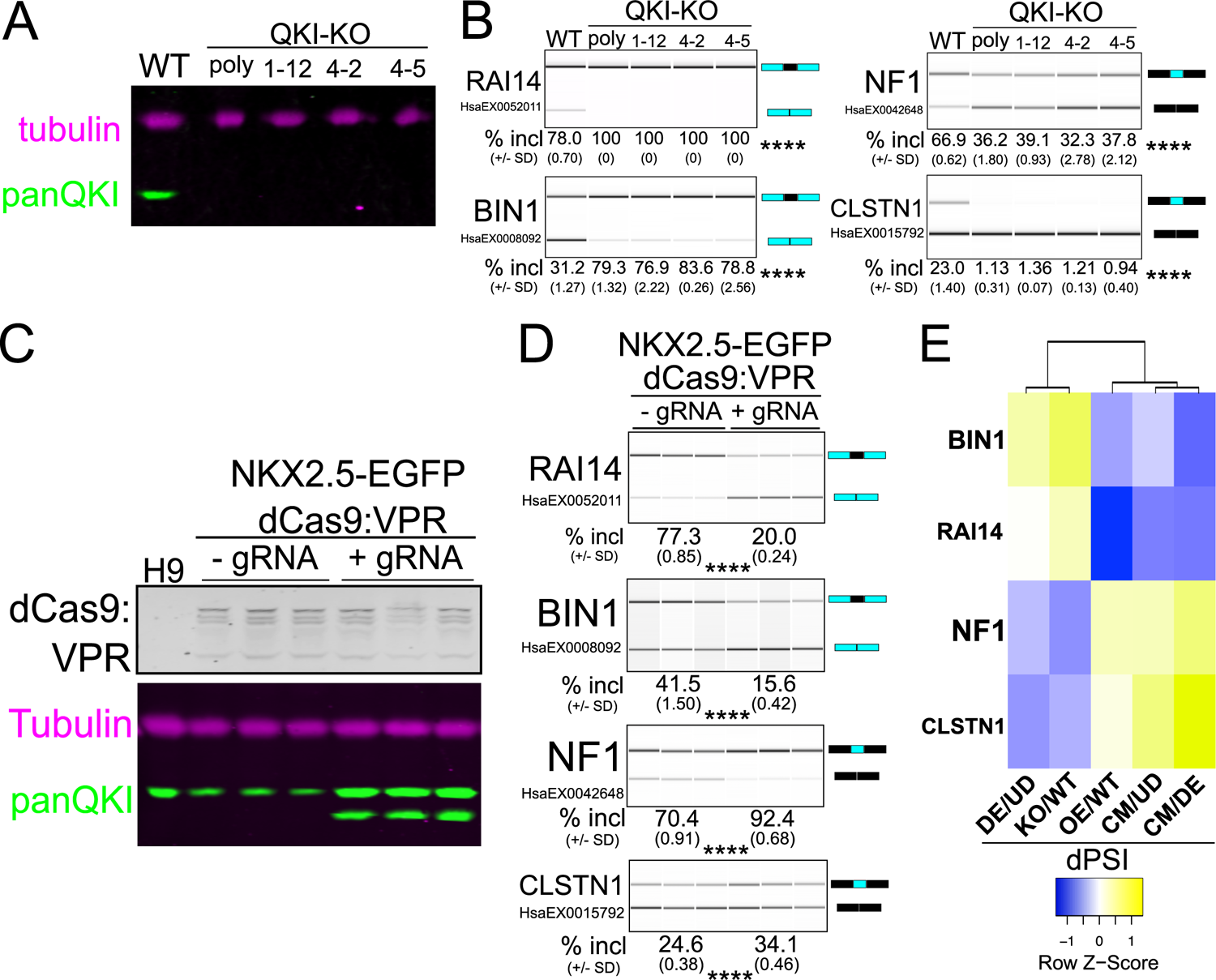
Loss of *QKI* results in changes in splicing of CM-activated exons. **A.** Western blot of protein extracted from UD *NKX2.5*➔EGFP WT, polyclonal *QKI*-KO (poly), 1-12 *QKI*-KO clone (1–12), 4-2 *QKI*-KO clone (4–2), and 4-5 *QKI*-KO clone (4–5) cells probed with antibodies for Tubulin (magenta) and panQKI antibodies (green; result shown is representative of 3 biological replicates). **B.** RT-PCR and BioAnalyzer gel-like image of RNA extracted from UD *NKX2.5*➔EGFP WT or *QKI*-KO cells described above to measure exon inclusion levels for *RAI14*, *BIN1*, *NF1*, and *CLSTN1* (result shown is representative of 3 biological replicates with mean percent included +/- standard deviation shown below; ****denotes *P* < 0.0001 for each KO cell extract relative to WT, by Student’s t-test). **C.** Western blot of protein extracted from UD H9 or triplicate cultured UD *NKX2.5*➔EGFP dCas9:VPR hESCs in the presence of doxycycline without (-gRNA) or with (+gRNA) QKI gRNA1, probed with anti-Cas9 antibody (top, gray), Tubulin (magenta), or panQKI (green). **D.** RT-PCR and BioAnalyzer gel-like image of RNA extracted from UD *NKX2.5*➔EGFP dCas9:VPR hESCs described above to measure alternative exon inclusion for *RAI14*, *BIN1*, *NF1*, and *CLSTN1* (****denotes *P* < 0.0001 for each +gRNA value relative to - gRNA value, by Student’s t-test). **E.** Clustered heatmap showing the correlation between dPSI values between DE cells relative to UD hESCs, UD *QKI*-KO hESCs relative to UD WT hESCs, UD hESCs with *QKI* overexpression relative to UD hESCs at basal levels, CM cells relative to UD cells, and CM cells relative to DE cells for *BIN1*, *RAI14*, *NF1*, and *CLSTN1* CEs calculated by RT-PCR and BioAnalyzer quantification and displayed as row Z-score values. See also Fig S3.

*RAI14* exon 11 and *BIN1* exon 7 cassette exons are skipped more in wild-type (WT) CM cells (Fig 1E), consistent with the prediction that they are QKI-repressed (Fig 2C), are more included in the QKI knock-out cells (Fig 3B). Conversely, *NF1* exon 23 and *CLSTN1* exon 11, which are more included in CM cells (Fig 1E), consistent with the prediction that they are QKI-activated (Figs 2C and S2B respectively), are more skipped in the QKI knock-out lines (Fig 3B). Also, *PICALM* exon 17, which is repressed in DE cells (Fig 1E), likely via QKI binding its downstream intron (Fig S2 B) and a reduction in QKI5 protein levels (Figs 2E), is less included in the absence of QKI (Fig S3 B). The CM cell-specific alternative exon observed in *DNMT3B* that was not predicted to be regulated by QKI is unaffected by QKI loss (Fig S3 C). These findings were independently validated following knockdown with two independent shRNAs targeting the QKI5 isoform (Figs S3D, S3E, S3F, and S3G).

We used a CRISPR activation (CRISPRa) approach (72) to increase the transcription of endogenous *QKI* (also in *NKX2.5*➔EGFP hESCs) and asked if this would produce opposite effects as those observed in the QKI-KO hESCs for the above splicing events. We confirmed transgene targeting to the *AAVS1* loci by genomic DNA PCR (data not shown), and doxycycline-inducible expression of dCas9:VPR and QKI protein isoforms by western blot (Fig 3C). A negative control consisting of H9 hESC lysate showed no reactivity with the Cas9 antibody, and the no guide RNA control *NKX2.5*➔EGFP+dCas9:VPR cell lysates showed basal levels of QKI protein (Fig 3C; compare to Fig S2 D also). Induction of *QKI* lead to a gain of function corresponding to the predicted splicing outcome: QKI-repressed events in *RAI14* and *BIN1* showed more skipping, QKI-activated events in *NF1* and *CLSTN1* showed more inclusion (Fig 3D), and the QKI-independent event in DNMT3B showed no change (Fig S3 H). Comparing the dPSI values for QKI-KO relative to WT hESCs, QKI overexpression (QKI-OE) relative to WT hESCs, CM cells relative to UD hESCs, DE cells relative to UD hESCs, and CM cells relative to DE cells for these events indicates a similar pattern for QKI-KO relative to WT hESCs and DE cells relative to UD hESCs, and also a distinct pattern in which CM cells compared to UD hESCs, CM cells compared to DE cells, and QKI-OE compared to WT hESCs are more similar (Fig 3E). Therefore, these CM cell-specific exons bound by QKI are responsive to both increases and decreases in QKI protein levels.

### QKI is required for efficient early cardiomyocyte differentiation

Our data provide evidence that QKI5 may directly promote up to 45% of CM lineage-specific CE splicing patterns (binding 59 out of 130 events; Fig S2 A and Table S4). We used the *QKI* null hESCs to compare the efficiency at which these and WT control cells form CM and early day 8 cardiomyocytes (8). QKI protein increases when WT hESCs differentiate to CM (Fig 2E) and cardiomyocytes (∼6-fold increase relative to tubulin), but as expected is undetectable in UD and day 8 polyclonal QKI-KO cells subjected to the cardiomyocyte differentiation protocol (Fig 4A). Interestingly, *QKI* KO does not affect *NKX2.5*➔EGFP levels during CM differentiation (Fig 4B) but the expression of this marker is significantly reduced upon further maturation to day 8 cardiomyocytes (Fig 4B and 4C; \**P* < 0.05, \*\**P* < 0.01 and \*\*\*\**P* < 0.001 respectively, by Student’s t-test), suggesting that QKI may be required for early cardiomyocyte differentiation.

**FIGURE 4:**
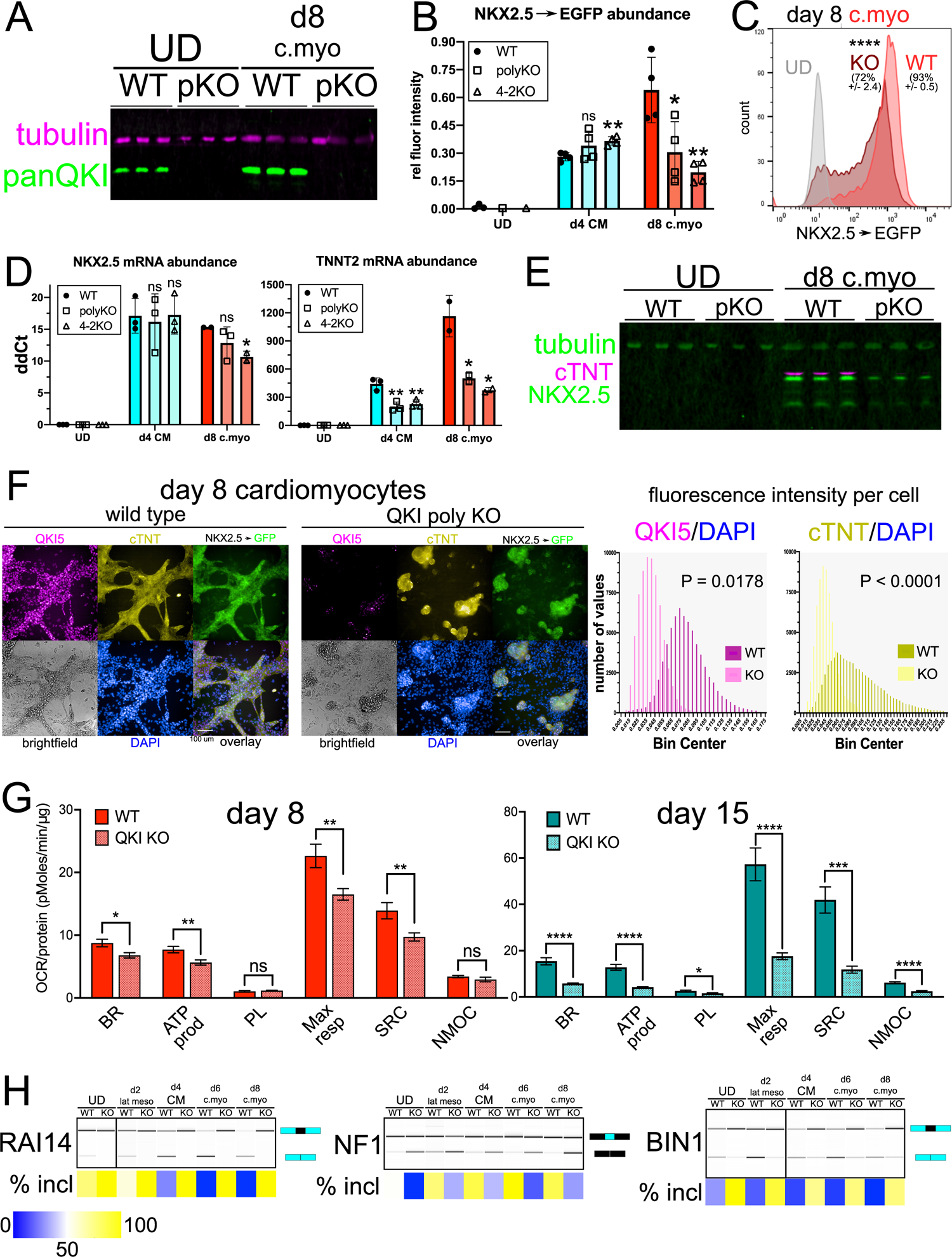
*QKI* is required for efficient cardiomyocyte cell differentiation. **A.** Western blot of protein extracted from *NKX2.5*➔GFP UD hESCs and day 8 cardiomyocyte (d8 c.myo) cells either wild-type (WT) or lacking *QKI* (polyclonal KO “pKO”) and probed with antibodies for tubulin (magenta) and panQKI (green). **B.** Population-level EGFP abundance (relative to GAPDH) calculated per well, measured by high content imaging in WT, polyclonal QKI-KO (polyKO), and clone 4-2 QKI-KO (4-2KO) in UD, day 4 CM (d4 CM), and day 8 cardiomyocytes (d8 c.myo) NKX2.5➔EGFP cells (n = 4 independent replicates; ns = not significant, \**P* < 0.05, and \*\**P* < 0.01 relative to WT, determined by Student’s t-test). **C.** Flow cytometry analysis of NKX2.5➔EGFP UD (gray), WT, or polyKO day 8 c.myo (light or dark red, respectively); mean percent GFP positive is shown above each histogram (+/- standard deviation; biological replicate n = 3) and \*\*\*\**P* < 0.0001 measured by Student’s t-test; y-axis denotes number of events measured. **D.** RT-qPCR measuring abundance of RNA extracted from UD hESCs, d4 CM cells, and d8 c.myo cells for endogenous *NKX2.5* and *TNNT2* (cTNT) mRNAs relative to *HMBS* mRNA in WT, polyKO, or 4-2KO cells (biological replicate n = 2 or 3 each; ns = not significant, \**P* < 0.05, \*\**P* < 0.01, measured by Student’s t-test). **E.** Western blot of protein extracted from *NKX2.5*➔EGFP UD hESCs and d8 c.myo cells either WT or pKO for *QKI* and probed with antibodies for tubulin (green, top) cTNT (magenta, middle (*TNNT2*)), and endogenous NKX2.5 (green, bottom). **F.** Indirect immunofluorescence with high content imaging analysis of *NKX2.5*➔EGFP WT (left) or QKI polyclonal KO (right) d8 cardiomyocyte cells showing QKI5 (magenta), cTNT (yellow), EGFP (green), brightfield (gray), and DAPI (blue) with each fluorescent channel overlaid (scale bar indicates 100 µm). On the right, the fluorescence intensity values per cell (relative to DAPI) are shown for QKI5 and cTNT with *P* values calculated by Mann–Whitney U test. **G.** Real-time mitochondrial bioenergetic measurement of intact day 8 or 15 cardiac differentiated WT or QKI-KO 4-2 cells measuring basal respiration (BR), ATP production (ATP prod), proton leak (PL), maximal respiration (Max resp), spare respiratory capacity (SRC), and non-mitochondrial O_2_ consumption (NMOC); the y-axes denote oxygen consumption rate (OCR) normalized to total protein content in pMoles/min/µg protein (n = 9 or 10 independent replicates each; ns = not significant, \**P* < 0.05, \*\**P* < 0.01, \*\*\**P* < 0.001, \*\*\*\**P* < 0.0001 by Student’s t-test). **H.** RT-PCR and BioAnalyzer gel-like image showing quantification of *RAI14*, *NF1*, and *BIN1* alternative exons using RNA extracted from WT or polyKO *NKX2.5*➔EGFP cells either UD, or subjected to the following differentiation protocols: day 2 lateral mesoderm (d2 lat meso), day 4 CM, day 6 cardiomyocyte (d6 c.myo), and day 8 cardiomyocyte (d8 c.myo) (biological replicate n=3; mean percent included +/- is shown below by heatmap; see Fig S4 G for mean numerical values and standard deviation from the mean). See also Fig S4.

To further investigate this, we measured endogenous biomarkers for CM and cardiomyocyte cell fate in *QKI*-KO and WT *NKX2.5*➔EGFP hESCs subjected to these differentiation protocols. Consistent with reporter EGFP levels, endogenous *NKX2.5* mRNA is unchanged in day 4 CM, but is significantly lower in the 4-2 clone KO day 8 cardiomyocyte cells (*P* < 0.05 by Student’s t-test; Fig 4D); cardiac Troponin T (cTNT; *TNNT2* gene), however, is lower in both CM and cardiomyocyte *QKI* KO cells (Fig 4D). Additional measurement of mRNA biomarkers for CM progenitor cells showed *ISL1* and *GATA4* abundance is lower in QKI KO cells, but *HAND1* and *MEF2C* do not significantly change (Fig S4 A). Thus, QKI loss may moderately disrupt CM cell differentiation or specifically perturb certain sub-populations of cardiac progenitor cells. We also used high content imaging to analyze these developmental transitions in WT and *QKI*-KO *NKX2.5*➔GFP hESCs, and observed no gross morphological perturbations, nor significant changes in OCT4 protein levels in the *QKI*-KO cells (Fig S4 B). However, on day 4 of differentiation to CM, endogenous NKX2.5 protein is significantly lower on a per-cell basis in the absence of QKI (*P* < 0.05 by Mann–Whitney U test; Fig S4 C) further supporting the notion of a moderate perturbation of CM differentiation in the absence of QKI. Interestingly however, by day 8, both NKX2.5 and cTNT protein levels are markedly lower in *QKI*-KO cells by western blot analysis (Fig 4E). High content imaging analysis (on both a per cell (Fig 4F) and population basis (Fig S4 D)) also indicate significantly reduced cTNT protein levels in day 8 *QKI*-KO cells (\*\*\*\**P* < 0.0001 by Mann–Whitney U and \*\*\**P* < 0.001 by Student’s t-test, respectively). Likewise, analysis of the morphology of *QKI*-KO cells indicates a reduction of branched cardiac muscle-like structures at day 8 (Fig 4F and S4E). *QKI* is required for late (day 15 and 30) hESC-derived cardiomyocyte cell contractile function and proper expression of structural and molecular markers (73), however it is an open question as to whether cardiomyocyte bioenergetic function is affected in the absence of QKI. Reduction of cardiac mitochondrial respiration can lead to embryonic or neonatal lethality, and cardiomyopathy (74–76), and thus is a well-established metric for measuring cardiomyocyte function (77). We measured this in both WT and clone 4-2 *QKI*-KO hESCs subject to day 8 or day 15 cardiomyocyte differentiation. Intriguingly, we observed significantly lower mitochondrial function in the absence of QKI in both day 8 and day 15 cardiomyocyte cells (Fig 4G). Specifically, basal respiration, ATP production-coupled respiration, maximal respiration, and spare respiratory capacity were all significantly reduced in *QKI*-KO hESCs at day 8 and to a greater magnitude at day 15 (upper bound \**P* < 0.05 by student’s t test; Fig 4F). Therefore, QKI is required for proper mitochondrial function during both earlier and later cardiomyocyte differentiation. We also tested if *RAI14* exon 11, *NF1* exon 23, and *BIN1* exon 7 inclusion levels were altered at various time points in these cells subjected to cardiac lineage differentiation. QKI is absolutely required for skipping of *RAI14* exon 11; *NF1* exon 23 inclusion increases during differentiation, but much less so in the absence of QKI; *BIN1* exon 7 skipping increases during differentiation, but is little changed in the absence of QKI (Figs 4H and S4F).Taken together, we conclude that QKI is required for proper cardiac differentiation.

### Complex regulation of BIN1 by QKI during cardiac cell lineage commitment

Interestingly, *Bin1* is required for cardiac development (78): specifically, either the absence or loss of expression of its cardiac-specific isoform results in improper calcium signaling or defective myofibril formation, respectively (79, 80). Given its importance in cardiac development and homeostasis, evidence of direct QKI binding (Fig S5 A), and its tissue-specific splicing patterns, we explored the consequences of QKI loss on its overall expression (Fig 5A). Accordingly, we cloned and sequenced full-length *BIN1* cDNAs from both UD and day 8 cardiomyocyte-differentiated WT and *QKI*-KO cells, and observed the expected increase in exon 7 inclusion in the absence of QKI, but also more inclusion of *BIN1* exon 13, the clathrin and AP (CLAP) binding domain-coding exons, and exon 17 (Figs S5A and 5A). Total *BIN1* mRNA levels in WT cells increases dramatically during cardiac differentiation. In the absence of QKI, the level is lower in UD hESCs, but slightly higher than WT in day 8 QKI-KO cardiomyocyte cells (Fig S5 B). Its protein levels in WT cells also increase during cardiac differentiation: a three-fold increase is observed by both western blot (Fig S5 C) and data-independent acquisition (DIA) LC-MS/MS (Fig S5 D and Table S7) when comparing UD hESCs to d8 cardiomyocyte cells. In QKI-KO cells, however, BIN1 protein levels are significantly lower in both CM (\**P* < 0.05 by Student’s t-test) and cardiomyocyte cells (\*\*\**P* < 0.001 by Student’s t-test; Fig S5 C). Moreover, the use of high-content imaging supported these findings when BIN1 protein was measured relative to GAPDH per well (Fig S5 E) and per cell in WT and QKI-KO day 8 cardiomyocytes (Fig S5 F). Therefore, during cardiac differentiation we observe complex changes in BIN1 expression in the absence of QKI.

**FIGURE 5:**
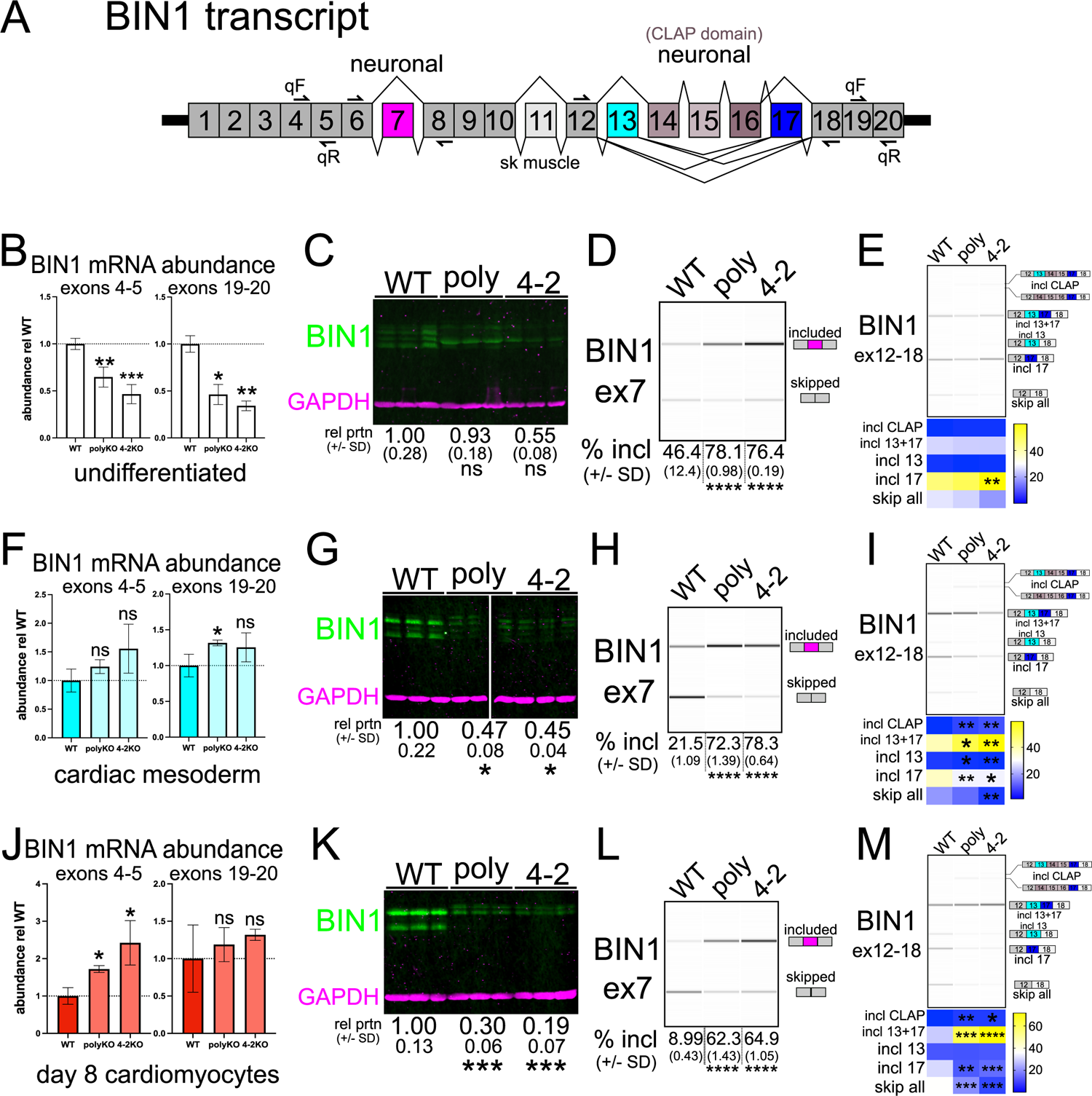
QKI promotes skipping of *BIN1* exon 7, exons 13+17, and exons 13-17 and total *BIN1* transcript maturation. **A.** *BIN1* transcript overview graphic: location of PCR and qPCR primers are noted, as are alternatively spliced exons and the tissue type with which they are correlated. **B.** RT-qPCR of RNA extracted from UD *NKX2.5*➔EGFP WT, polyKO, or 4-2KO hESCs measuring either the 5’ end of *BIN1* (primer pair spanning exons 4 and 5) or the 3’ end of *BIN1* (primer pair spanning exons 19 and 20); value shown is ddCt relative to HMBS and normalized to WT; n = 2 or 3 independent replicates, ns = not significant, \**P* < 0.05, \*\**P* < 0.01, \*\*\**P* < 0.001 by Student’s t-test compared to WT. **C.** Western blot of protein extracted from UD WT, polyKO, and 4-2KO *NKX2.5*➔EGFP hESCs probed with a BIN1 antibody that recognizes all isoforms (green) and GAPDH antibody (magenta); three independent biological replicates are shown, with mean protein levels relative to WT and normalized to GAPDH shown below +/- standard deviation from the mean (ns denotes not significant by Student’s t-test). **D.** RT-PCR and BioAnalyzer gel-like image showing quantification of *BIN1* exon 7 inclusion using RNA extracted from UD WT, polyKO, or 4-2KO NKX2.5➔EGFP hESCs (biological replicates of n=3; mean percent included values +/- standard deviation are shown below; \*\*\*\**P* < 0.0001 by student’s t test). **E.** RT-PCR and BioAnalyzer gel-like image showing quantification of *BIN1* products amplified by primers annealing to exons 12 (forward) and 18 (reverse) using RNA extracted from UD WT, polyKO, or 4-2KO *NKX2.5*➔EGFP hESCs (biological replicate n=3; mean percentage of product detected is shown below, represented by intensity plot; \*\**P* < 0.01 by student’s t test). **F.** RT-qPCR of RNA extracted from d4 CM *NKX2.5*➔EGFP WT, polyKO, or 4-2KO cells measuring either the 5’ or 3’ end of *BIN1* as in Fig 5B; n = 2 or 3 independent replicates, ns = not significant, \**P* < 0.05 by Student’s t-test compared to WT. **G.** Western blot of protein extracted from d4 CM WT, polyKO, and 4-2KO *NKX2.5*➔EGFP cells probed with BIN1 and GAPDH antibodies as described in Fig 5C; \**P* < 0.05 by Student’s t-test. **H.** RT-PCR and BioAnalyzer gel-like image showing quantification of *BIN1* exon 7 inclusion using RNA extracted from d4 CM WT, polyKO, or 4-2KO *NKX2.5*➔EGFP cells as in Fig 5D. **I.** RT-PCR and BioAnalyzer gel-like image showing quantification of *BIN1* products amplified by primers annealing to exons 12 (forward) and 18 (reverse) using RNA extracted from d4 CM WT, polyKO, or 4-2KO *NKX2.5*➔EGFP cells as in Fig 5E (\**P* < 0.05, \*\**P* < 0.01 by student’s t test). **J.** RT-qPCR of RNA extracted from d8 cardiomyocyte *NKX2.5*➔EGFP WT, polyKO, or 4-2KO cells measuring either the 5’ or 3’ end of *BIN1* as in Fig 5B; n = 2 or 3 independent replicates, ns = not significant, \**P* < 0.05 by Student’s t-test compared to WT. **K.** Western blot of protein extracted from d8 cardiomyocyte WT, polyKO, and 4-2KO *NKX2.5*➔EGFP cells probed with BIN1 and GAPDH antibodies as described in Fig 5C; \*\*\**P* < 0.001 by Student’s t-test. **L.** RT-PCR and BioAnalyzer gel-like image showing quantification of *BIN1* exon 7 inclusion using RNA extracted from d8 cardiomyocyte WT, polyKO, or 4-2KO *NKX2.5*➔EGFP cells as in Fig 5D. **M.** RT-PCR and BioAnalyzer gel-like image showing quantification of *BIN1* products amplified by primers annealing to exons 12 (forward) and 18 (reverse) using RNA extracted from d8 cardiomyocyte WT, polyKO, or 4-2KO *NKX2.5*➔EGFP cells as in Fig 5E (\**P* < 0.05, \*\**P* < 0.01, \*\*\**P* < 0.001, \*\*\*\**P* < 0.0001 by student’s t test).

To specifically determine how QKI regulates *BIN1* transcript maturation during cardiac differentiation, we measured RNA and protein abundance, and splicing of exon 7 and the region encompassing exons 12 through 18 in WT and QKI-KO UD hESCs, CM cells, and day 8 cardiomyocyte cells. In UD QKI-KO hESCs, *BIN1* mRNA is significantly reduced (upper bound \**P* < 0.05 by Student’s t-test, see Fig 5B). Its protein levels do not significantly change, but there is a clear shift to a prominent band of a lower molecular weight (Figs 5C and S5C). It is unknown if this represents an isoform change or gain/loss of a post-translational modification. As measured previously (Figs 3B and 4G), exon 7 is significantly more included in the absence of QKI (\*\*\*\**P* < 0.0001 by Student’s t-test; Fig 5D), and we observed more inclusion of exon 17 in the 4-2KO clonal cell line (Figs 5E and S5G). We conclude that QKI is required for maintaining *BIN1* transcript levels and exon 7 skipping in UD hESCs.

In QKI-KO cells subject to CM differentiation, there is a slight increase in *BIN1* mRNA compared to WT (Fig 5F), but a significant reduction in its protein levels (\**P* < 0.05 by Student’s t-test; Fig 5G). We also observe more inclusion of exon 7 in *QKI*-KO CM cells (Fig 5H), and a more dramatic change in its downstream exons: the CLAP domain exons and exons 13+17 are included more in KO, and exon 17 is skipped more (Figs 5I and S5G). Therefore, in CM cells, QKI is required for BIN1 protein accumulation and proper splicing, but total *BIN1* mRNA levels are unchanged in the absence of QKI.

We measured the 5’ end of *BIN1* mRNA and found a significant increase in its level (\**P* < 0.05 by Student’s t-test) in *QKI*-KO cells subject to day 8 cardiomyocyte differentiation, but this was not apparent when we measured the 3’ end (Fig 5J). In the absence of QKI, we observed large magnitude reductions in BIN1 protein levels by western blotting (***P < 0.001 by Student’s t-test; Fig 5K). Consistent with the above results (Fig 4G), exon 7 is included more in the absence of QKI (Fig 5L). Finally, the downstream alternative exons are more impacted in *QKI*-KO cardiomyocyte cells than in UD or CM cells: inclusion of CLAP domain-encoding exons increased, inclusion of exons 13 + 17 are significantly higher (\*\*\**P* < 0.001 or \*\*\*\**P*< 0.0001 by Student’s t-test for polyKO or 4-2KO, respectively) with a concomitant increase in exon 17 skipping and the “skip all” isoform (Figs 5M and S5G). It is unlikely that mis-processing of BIN1 alone is sufficient for the early human cardiomyocyte defects we observed in *QKI*-KO cells (Fig 4), since QKI loss results in over 1000 transcript-level and 461 splicing changes in *QKI*-KO day 15 cardiomyocytes (73). However, our data do indicate a loss of “muscle-specific” (81) and a gain of “neuron-specific” *BIN1* isoforms (82–85), in the absence of QKI (see also Discussion). Taken together, we conclude that QKI is required for cardiac muscle-specific BIN1 transcript processing and protein accumulation.

### QKI promotes BIN1 exon 7 skipping and nuclear export

To explore the potential molecular mechanisms through which QKI regulates *BIN1* exon 7 splicing, we generated a splicing reporter containing this exon, along with 126 bp of upstream and 184 bp of downstream intron sequence (41). Transfection of this reporter into HuH7 WT or *QKI*-KO cells (39, 86) and analysis by RT-PCR showed a modest increase in exon 7 inclusion in the absence of QKI (Fig 6A), consistent with QKI promoting skipping of this exon. Strikingly, the largest magnitude changes observed were an increase in unspliced reporter RNA and a decrease in exon 7 skipping (Fig 6A). Transfection of HuH7 *QKI*-KO cells with myc:Qk5 fully repressed inclusion, and partially rescued unspliced RNA accumulation, despite its expression level being below that observed in WT HuH7 cells (Figs 6A and 6B). Interestingly, adding back an RNA-binding mutant version of Qk5 (K120A/R124A (87)) did not affect reporter splicing by a large magnitude, compared to vector-transfected control *QKI*-KO HuH7 cells (Figs 6A and 6B). We conclude that *QKI* is necessary and sufficient to promote BIN1 exon 7 skipping and reduce accumulation of unspliced reporter RNA.

**FIGURE 6:**
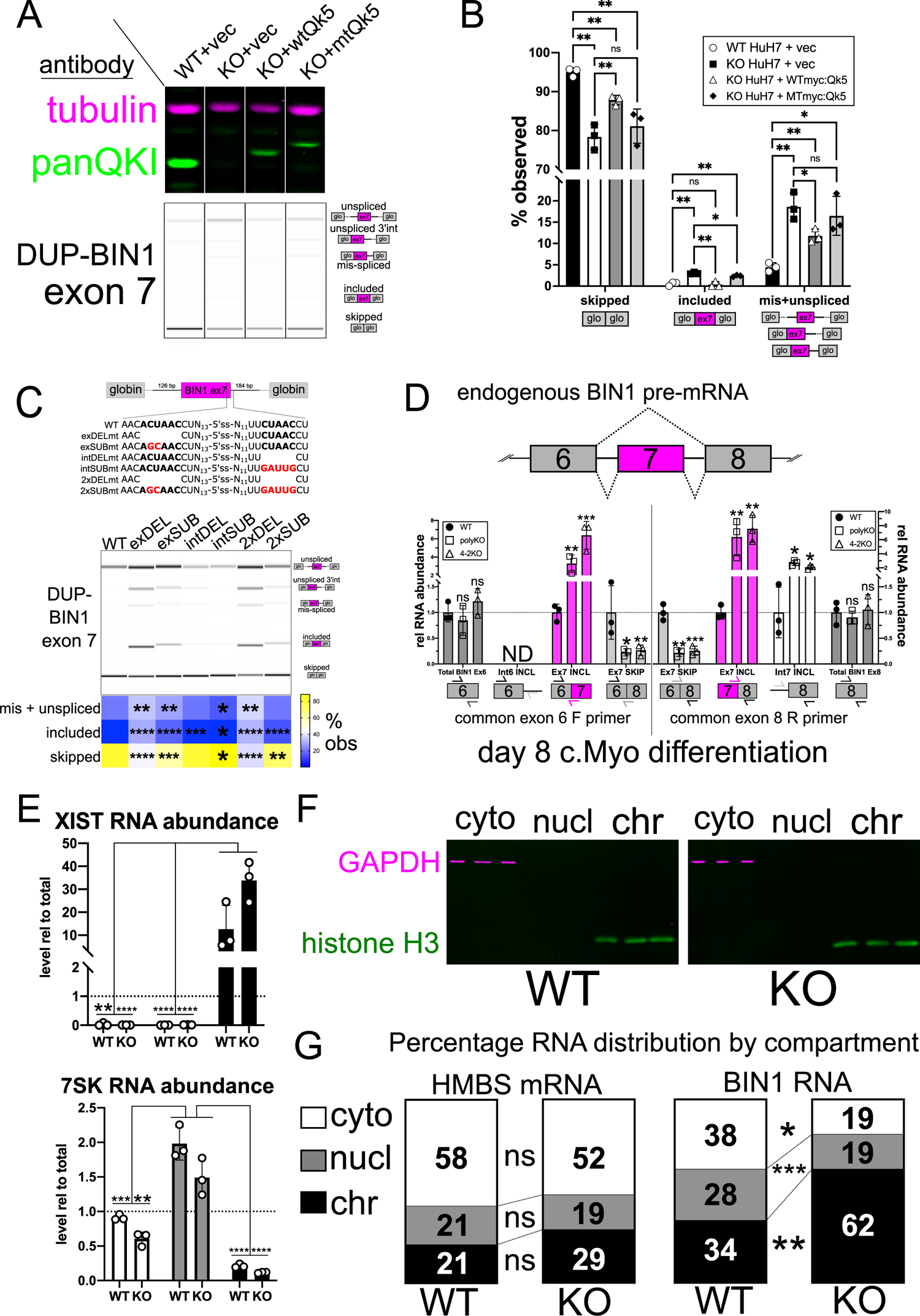
QKI is required for efficient BIN1 pre-mRNA processing. **A.** Representative western blot (top) of protein extracted from three independent replicates of HuH7 WT or HuH7 *QKI*-KO cells co-transfected with pDUP-BIN1 exon 7 splicing reporter and either vector control (vec), myc:Qk5 (wtQk5), or K120A/R124A myc:Qk5 (mtQk5) probed with panQKI antibody (green) and Tubulin antibody (magenta). Representative RT-PCR and BioAnalyzer gel-like (bottom) image showing splicing reporter products derived from RNA extracted from WT or *QKI*-KO cells described above. **B.** Bar graph showing mean percentage of each RT-PCR product (or products) observed in Fig 6A, under each experimental condition described (error bars denote standard deviation from the mean; ns denotes not significant, *P < 0.05, **P < 0.01; note that in the absence of minus reverse transcription controls we cannot exclude the possibility that reporter DNA might have been amplified and co-migrate with “unspliced”). **C.** Overview graphic of pDUP-BIN1 exon 7 splicing reporter (top) with WT and mutant sequences shown in inset below. Representative RT-PCR and BioAnalyzer gel-like image (bottom) showing quantification of WT or mutant splicing reporter products derived from RNA extracts from WT HuH7 cells transfected in triplicate; the mean percentage observed of each product(s) is shown below the gel-like image represented by intensity plot (\**P* < 0.05, \*\**P* < 0.01, \*\*\**P* < 0.001, \*\*\*\**P* < 0.0001 by student’s t test; see also Fig S6 B for mean values +/- standard deviation and minus reverse transcription controls). **D.** RT-qPCR of RNA extracted from d8 c.myo *NKX2.5*➔EGFP WT, polyKO, or 4-2KO cells measuring (from left to right) total exon 6 (“total”; primers internal to exon 6 including common exon 6 forward), intron 6 included (“int6+”; common forward primer in exon 6 with reverse primer in intron 6), exon 6-7 spliced (“ex7+”; common forward primer in exon 6 with reverse primer spanning exon 6:7 junction), exon 6-8 spliced (“ex7-“; common forward primer in exon 6 with reverse primer spanning exon 6:8 junction), exon 6-8 spliced (“ex7-“; forward primer spanning exon 6:8 junction with common reverse primer in exon 8) exon 7-8 spliced (“ex7+”; forward primer spanning exon 7:8 junction with common reverse primer in exon 8), intron 7 included (int7+; forward primer in intron 7 with the common reverse primer in exon 8), and total exon 8 levels (“total”; forward primer in exon 8 with the common reverse primer in exon 8) from the endogenous *BIN1* transcript; each value is shown normalized to *HMBS* mRNA and relative to WT; graphic below bars indicates approximate location of forward and reverse primer pairs; n = 3 biological replicates, ns = not significant, \**P* < 0.05, \*\**P* < 0.01, \*\*\**P* < 0.001, relative to WT by Student’s t-test). **E.** RT-qPCR of RNA extracted from d8 c.myo NKX2.5➔EGFP WT or polyKO cells’ cytoplasmic (cyto), nucleoplasmic (nucl), or chromatin (chr) fractions, measuring *XIST* (top) or *7SK* (bottom) RNAs. Their levels are shown normalized to *HMBS* mRNA and relative to total (unfractionated) cell extract RNA levels (\*\**P* < 0.01, \*\*\**P* < 0.001, \*\*\*\**P* < 0.0001 by student’s t test). **F.** Western blot of protein extracted from subcellular fractions of WT (left) or polyKO (right) d8 c.myo *NKX2.5*➔EGFP cells as described above and probed with anti-GAPDH (magenta) and anti-histone H3 (green) antibodies. **G.** RT-qPCR of RNA extracted from subcellular fractions of WT or polyKO d8 c.myo *NKX2.5*➔EGFP cells as described above and measuring *HMBS* (left) or *BIN1* (right) RNA, shown as the percentage of RNA observed by compartment, relative to total RNA from unfractionated extracts; ns denotes not significant, *P < 0.05, **P < 0.01, ***P < 0.001 by student’s t test.

Closer inspection of *BIN1* exon 7 and surrounding sequences revealed a potential exonic QKI motif ACUAAC and putative “half-site” in the downstream intron CUAAC (Fig 6C). Analysis of iCLIP reads from C2C12 myoblasts using the panQk antibody (33) indicated direct binding to this region (Fig S6 A). We hypothesized that these motifs are required for proper splicing, and tested this by generating exonic or intronic deletion or substitution mutants, and double mutants to ablate the respective motifs in the reporter then measured their splicing in WT HuH7 cells (Fig 6C). Overall, deletion or substitution of the exonic motif resulted in more inclusion of exon 7, unspliced RNA, and mis-spliced RNA; the intronic mutants alone affected little to no, or very small magnitude changes (Figs 6C and S6B). We next asked whether QKI levels impact both exon 7 skipping and intron retention of endogenous *BIN1* transcripts using *QKI*-KO hESCs subjected to 8 day cardiomyocyte differentiation. We measured total *BIN1* RNA levels (exon 6 or exon 8 internal primers), intron 6, exon 7, and intron 7 inclusion by RT-qPCR (see schematic and figure legend for more information; Fig 6D). Importantly, the concomitant increases in both exon and intron 7 inclusion observed in the reporter assays were also apparent in endogenous *BIN1* transcripts in the absence of QKI (Fig 6D). Therefore, QKI and the exonic motif in *BIN1* exon 7 promote exon skipping and intron removal.

We hypothesized that un- or mis-processed *BIN1* RNA might be retained on chromatin in the absence of QKI, which could explain the increase in total RNA accumulation but the observed sharp reduction in BIN1 protein. To test this, we subjected WT or *QKI*-KO hESCs to an 8 day cardiomyocyte differentiation protocol, performed subcellular fractionation, and measured protein and RNA levels. The chromatin-associated lncRNA *XIST* was significantly enriched in the chromatin fraction (\*\*\*\**P* < 0.0001 or \*\**P* < 0.01 by Student’s t-test), and the nucleoplasmic RNA *7SK* was enriched in the nucleoplasmic fraction (Fig 6E). Western blotting analysis indicated the presence of GAPDH within the cytoplasmic fraction, and histone H3 in the chromatin fraction (Fig 6F), therefore fraction-specific components are enriched in their cognate compartments. Next, we measured a transcript not predicted to be directly regulated by QKI (*HMBS*) in WT and KO cells, and observed no significant difference between their distributions (Fig 6G). In contrast, total *BIN1* RNA is significantly enriched in the chromatin fraction and depleted from the nucleoplasmic and cytoplasmic fractions (upper bound is *P* ≤ 0.05 by Student’s t-test; Fig 6G). These observations support a model in which QKI promotes the processing of *BIN1* transcripts to facilitate chromatin release and subsequent nucleocytoplasmic export, thereby facilitating BIN1-dependent functions during cardiogenesis.

## DISCUSSION

Studies in model organisms indicate *Quaking* is required for embryonic development (26,88,89), and particularly so in mesoderm/muscle formation (89–94) and neural cell specification (88, 95). However, its roles at the earliest stages of lineage specification and cardiomyocyte formation had not previously been investigated. Here we uncovered that this RBP significantly increases or decreases during the exit from pluripotency and commitment to either mesoderm or endoderm, respectively. We also found extensive evidence of QKI direct binding to, and motif enrichment in and around, cardiac lineage-specific alternatively spliced exons. Additionally, we observed many differentiation- and lineage-specific changes in the levels of other RBPs, and certainly there are important regulatory interactions between some of these and QKI. However, given the significant representation of QKI described above, we propose that it is a mesoderm-enriched splicing regulator that critically shapes the transcriptome during lineage-commitment.

We also find that QKI is required for efficient early cardiomyocyte differentiation, and regulates multiple exons in genes linked to cardiac tissue development (78, 96). Among these exons we focused on the QKI-dependent repression of exon 7 of BIN1. Interestingly, our data suggest that QKI regulates *BIN1* splicing through a set of unique molecular mechanisms, in which QKI binds to an exonic sequence motif to coordinately promote skipping of exon 7, removal of intron 7, and then ligation of exons 6 and 8, which licenses nuclear *BIN1* mRNA export; these regulatory events are required for BIN1 protein accumulation in hESC-derived cardiomyocytes (Figs 5 and 6). An exonic ACUAA motif is critical for exon 7 skipping, downstream intron removal, and proper 3’ splice site usage in our splicing reporter (Fig 6), suggesting that characterizing this motif simply as a “splicing silencer” is inadequate. QKI and this exonic motif cooperate to ensure proper quantitative and qualitative expression of *BIN1* in human cardiomyocytes, which is required for their differentiation and homeostasis. We do note it is unlikely that rescuing *BIN1* expression alone in QKI-KO cells would be sufficient to restore cardiomyocyte differentiation, as QKI regulates thousands of transcripts during this process (73). Furthermore the *Bin1* knockdown cardiac phenotype is more nuanced (80) than the one we have observed here, in *qki^-/-^* (89) and in other *qk* mutant mice (97). However, these *BIN1*-specific regulatory events are critically important: inclusion of BIN1 exon 7 in skeletal muscle is associated with centronuclear myopathy in myotonic dystrophy patients (98), and functions in controlling endocytosis via promoting an interaction with Dnm2 in neural cells, where it is positively regulated by SRRM4 (82). In post mitotic neurons, QKI is absent (95). Other *BIN1* exons, 13, 13-16 (CLAP domain), and exon 17, included in neurons but skipped in cardiomyocytes, are also repressed by QKI (Fig 5), suggesting that cardiac versus neuronal *BIN1* splicing patterns are determined by QKI. Dysregulation of total *BIN1* levels and splicing of these exons in the brain is correlated with Alzheimer’s disease (99, 100). Therefore, the tight regulation of *BIN1* levels and isoform output is required for proper tissue homeostasis and differentiation in cardiac muscle, skeletal muscle, and the brain, and we establish that QKI directly regulates these via RNA splicing.

The gene expression process is highly regulated, and subject to dynamic on/off states as well as finely tuned rheostat-like adjustments to properly respond to various inputs (101–103). Extensive mapping of chromatin/histone and promoter states, and transcription factor and RNA polymerase II occupancy during commitment of hESCs to each of the three germ layers has identified both unique and shared states between these different progenitor cell types (11, 13). Consistent with previous findings (13), we observe quantitative changes in gene expression that are shared between the three germ layer derivatives (Fig S1 D). This implies that most transcriptional on/off states are developmentally regulated, and are not lineage-specific. In turn, these are targets of alternative splicing, which can have profound effects on the ultimate gene product repertoire. Interestingly, the splicing patterns of DE and CM cells are uniquely different from hESCs, and even more different from one another (Fig 1). Of particular interest is that several exons in genes coding for transcription/chromatin modifying factors that promote differentiation are markedly different between DE and CM cells. These resemble examples of splicing level regulation, the products of which can profoundly alter the transcriptional response, either directly (i.e., splicing of the transcription factor *FOXP1*, which can control transitions between pluripotency and differentiation (60)) or indirectly by altering chromatin states (i.e., splicing of the DNA methylase *DNMT3B*, which can control DNA methyltransferase activity and gene body methylation (62, 63)). Therefore, modulation of gene expression via alternative splicing can influence downstream transcriptional regulatory events in a lineage-specific manner. While certainly not all of these splicing changes can be attributed to QKI, we find that about 45% of them in the cardiac lineage are, implicating QKI as one possible master regulator splicing factor (25). Orchestration of these complex and oftentimes combinatorial molecular-level transitions by QKI is critical for the exit from pluripotency and commitment to lineage-specific gene expression.

## Supporting information

Oligonucleotide sequences are listed in Supplementary Table S8.

## ACCESSION NUMBERS

The RNAseq data was deposited at Gene Expression Omnibus (GSE162649); proteomics datasets were deposited at MassIVE (MSV000088762).

## SUPPLEMENTARY DATA

Supplementary data are available at NAR online.

## ACKNOWLEDGEMENTS

We thank Rhonda Perriman for pre-submission review and valuable comments. We thank Seigo Hatada, David Elliot, Andrew Elefanty, and Ed Stanley for hESCs, and Kuo-Chieh Liao for HuH7 cells and QKI sgRNA vector.

## AUTHOR’S CONTRIBUTIONS

Conceptualization, WSF.; Methodology, WSF; Formal analysis, WSF, UB, XC, JPD, WKR, SGW, MTW; Investigation, WSF, NL, UB, KLPC, KM, WKR, FSD, XC; Resources, WSF, JHF, MTW, BJB, MAGB; Writing – Original Draft, WSF, Writing – Review and Editing, WSF, UB, SGW, MTW, BJB, MAGB; Visualization, WSF, UB; Supervision WSF; Project Administration WSF; Funding Acquisition WSF, JHF, KM, WKR, MTW, BJB, MAGB.

---

**SUPPLEMENTAL FIGURE 1:**
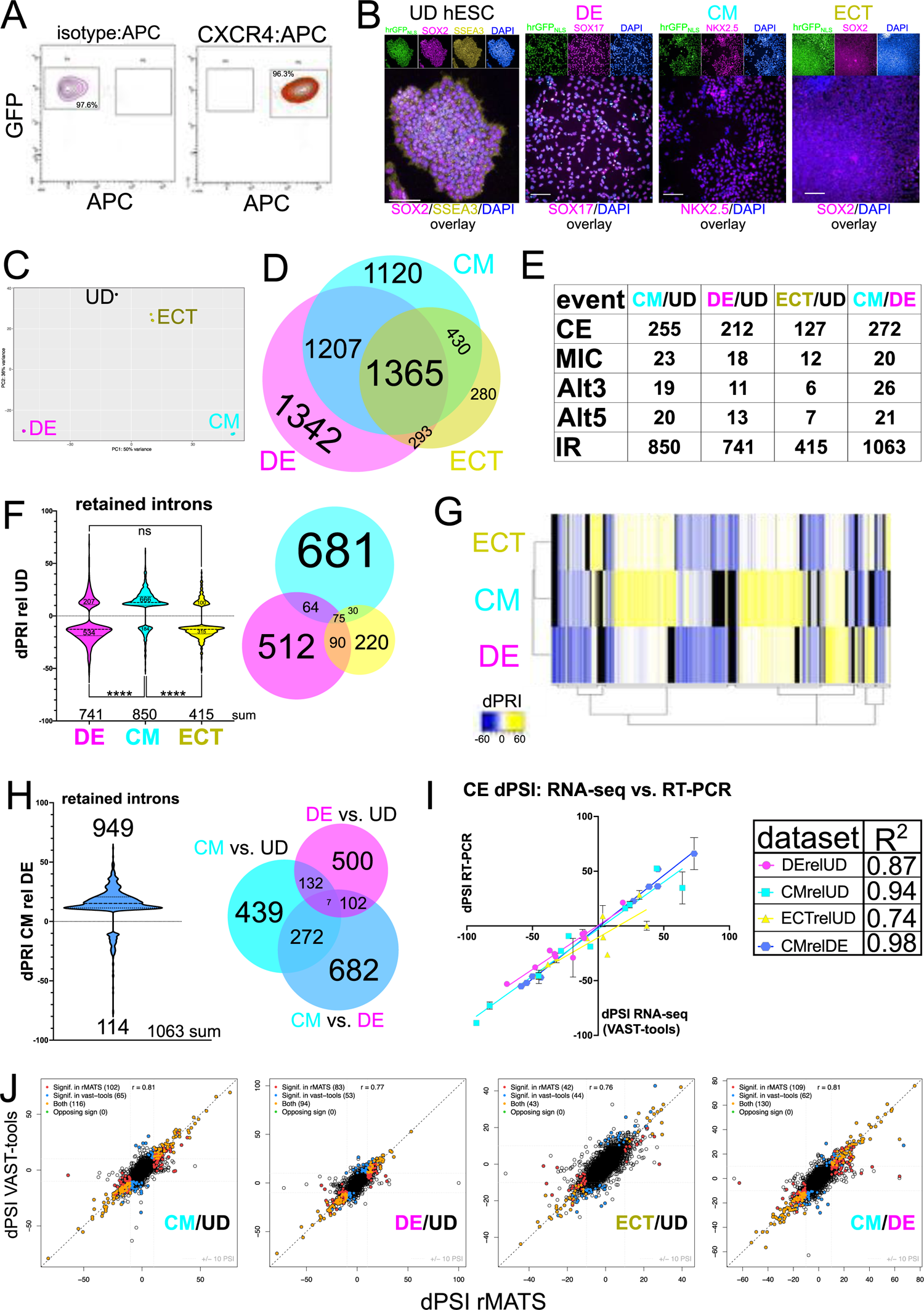
A. Fluorescence-activated cell sorting (FACS) strategy showing density plots for day 3 differentiated DE cells’ antibody isotype control (right) and a single representative of three independent replicates for CXCR4:APC stained cells (right) that were selected by FACS based on GFP (y-axis) and CXCR4 (x-axis) dual positivity and from which RNA was extracted for RNA-seq (the cells constitutively express a nuclear localized humanized renilla GFP (hrGFP-NLS)). B. Indirect immunofluorescence of H9 hESCs with hrGFP-NLS, showing hrGFP (green), SOX2 (magenta), SSEA3 (yellow), and DAPI (blue; SOX2, SSEA3, and DAPI in overlay) for UD hESCs; hrGFP (green), SOX17 (magenta), and DAPI (blue; SOX17 and DAPI overlay) for DE cells, hrGFP (green); NKX2.5 (magenta), and DAPI (blue; NKX2.5 and DAPI overlay) for CM cells; and hrGFP (green), SOX2 (magenta), and DAPI (blue; SOX2 and DAPI overlay) for ECT cells (scale bar = 100 μm). C. Principal component analysis plot of the top 500 genes with the most variance between replicates of UD, DE, CM, and ECT cells, clustered by similarity (n = 3 each). D. Overlap of protein-coding transcript abundances that significantly changed in CM, DE, and ECT cells relative to UD hESCs from DESEQ2 analysis of RNA-seq data. E. Table showing total number of alternatively spliced events significantly changing in CM cells relative to UD (CM/UD), DE cells relative to UD (DE/UD), ECT cells relative to UD (ECT/UD), or CM cells relative to DE cells (CM/DE) by VAST-tools analysis of RNA-seq data. F. Change in percent retained intron (dPRI) for intron retention events (positive values indicate an increase in intron retention, lower values indicate a decrease) in DE, CM, and ECT cells, significantly changing relative to UD hESCs in violin plot, and overlap of these shown in Venn diagram inset (ns = not significant, and \*\*\*\**P* ≤ 0.001 by Mann-Whitney test). G. Heatmap showing hierarchical clustering of significantly changing retained introns from DE, CM, and ECT cells, calculated by dPRI relative to UD hESCs. H. Comparison of dPSI values calculated by RT-PCR and BioAnalyzer quantitation (y-axis) versus VAST-tools analysis of RNA-seq (x-axis) data for each alternatively spliced exon shown in Fig 1E; R^2^ values are shown for DE cells compared to UD hESCs (DErelUD), CM cells compared to UD hESCs (CMrelUD), ECT cells compared to UD hESCs (ECTrelUD), and CM cells compared to DE cells (CMrelDE) in the inset. I. dPRI of significantly changing retained introns for CM cells relative to DE cells shown in violin plot, and the overlap of these compared to CM cells relative to UD hESCs and DE cells relative to UD hESCs shown in the Venn diagram inset, calculated from VAST-tools analysis of RNA-seq data. J. Dot plot comparing dPSI values obtained by VAST-tools (y-axis) to dPSI values obtained by rMATS for CE/MIC events indicating those significant in rMATS (red), VAST-tools (blue), both (orange), or significant in both but in opposite directions (Opposing sign; green), or neither (open circle) for CM/UD, DE/UD, ECT/UD, or CM/DE.

**SUPPLEMENTAL FIGURE 2:**
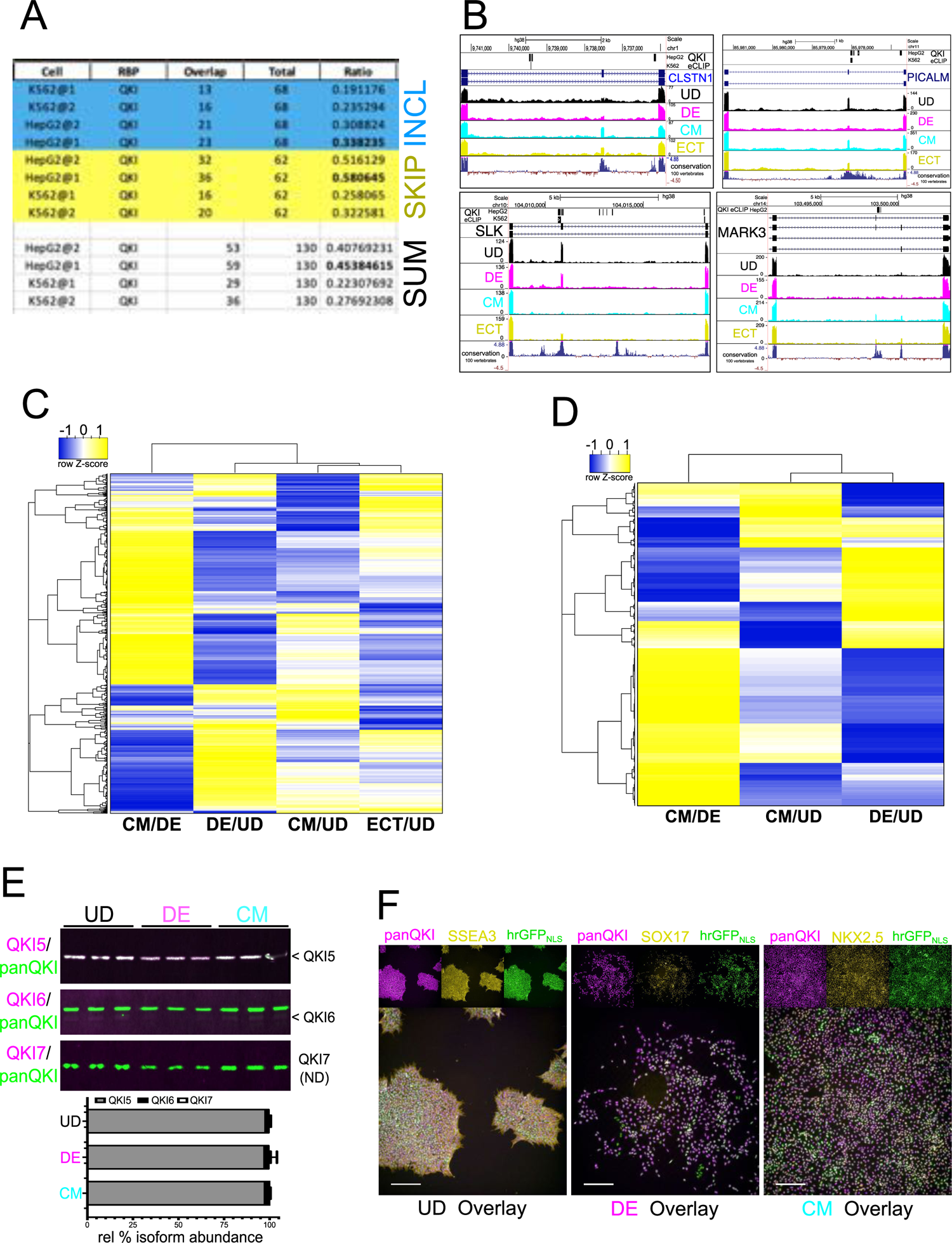
A. Summary of QKI binding frequency to lineage-specific alternatively spliced exons identified by both VAST-tools and rMATS analysis. “Cell” denotes cell type in which CLIP experiment was performed (@1 refers to experimental replicate number one; @2 refers to experimental replicate number two), “RBP” is the RBP of interest (QKI), “Overlap” is the number of occurrences in which merged coordinates of interest (upstream intron, alternatively spliced exon, and downstream intron) intersect with at least one QKI CLIP peak, “Total” is the total number of cassette exons that passed significance cutoff in both VAST-tools and rMATS, and “Ratio” is the ratio of overlap to total (values in bold are the highest observed for all “Cell” conditions), which measures the ratio of QKI binding to cassette exons and flanking intron sequence. CEs that are more included in CM cells relative to DE cells are shown in blue, more skipped CEs are shown in yellow, with the sum of events below. B. UCSC Genome Browser screen shots showing QKI eCLIP peaks, Gencode Gene annotation for *CLSTN1, PICALM, SLK,* and *MARK3*, with RNA-seq coverage tracks for UD hESCs (black), DE cells (magenta), CM cells (cyan), and ECT cells (yellow), and conservation of 100 vertebrates track. C. Heatmap showing hierarchical clustering of annotated RBPs that change significantly in at least one lineage by DESEQ2 analysis of RNA-seq data; log_2_ fold change values from CM cells relative to DE cells, DE cells relative to UD hESCs, CM cells relative to UD hESCs, and ECT cells relative to UD hESCs were used to make the heatmap and are displayed as row Z-score values. D. Heatmap showing hierarchical clustering of annotated RBPs that change significantly in at least one lineage by LC-MS/MS analysis; log_2_ fold change values from CM cells relative to DE cells, CM cells relative to UD hESCs, and DE cells relative to UD hESCs were used to make the heatmap and are displayed as row Z-score values. E. Western blot of protein extracted from UD hESCs, DE cells, and CM cells probed with antibodies for QKI5 (magenta, top panel) and panQKI (green), QKI6 (magenta, middle panel) and panQKI (green), or QKI7 (magenta, bottom panel; ND denotes not detectable) and panQKI (green); quantitation of each isoform relative to total panQKI is shown below by bar graph. F. Indirect immunofluorescence of H9 hrGFP-NLS showing panQKI (magenta), SSEA3 (yellow), hrGFP (green), and each overlaid for UD hESCs; panQKI (magenta), SOX17 (yellow), and hrGFP (green), with each overlaid for DE cells; panQKI (magenta), NKX2.5 (yellow), and hrGFP (green), with each overlaid for CM cells (scalebar denotes 100µm).

**SUPPLEMENTAL FIGURE 3:**
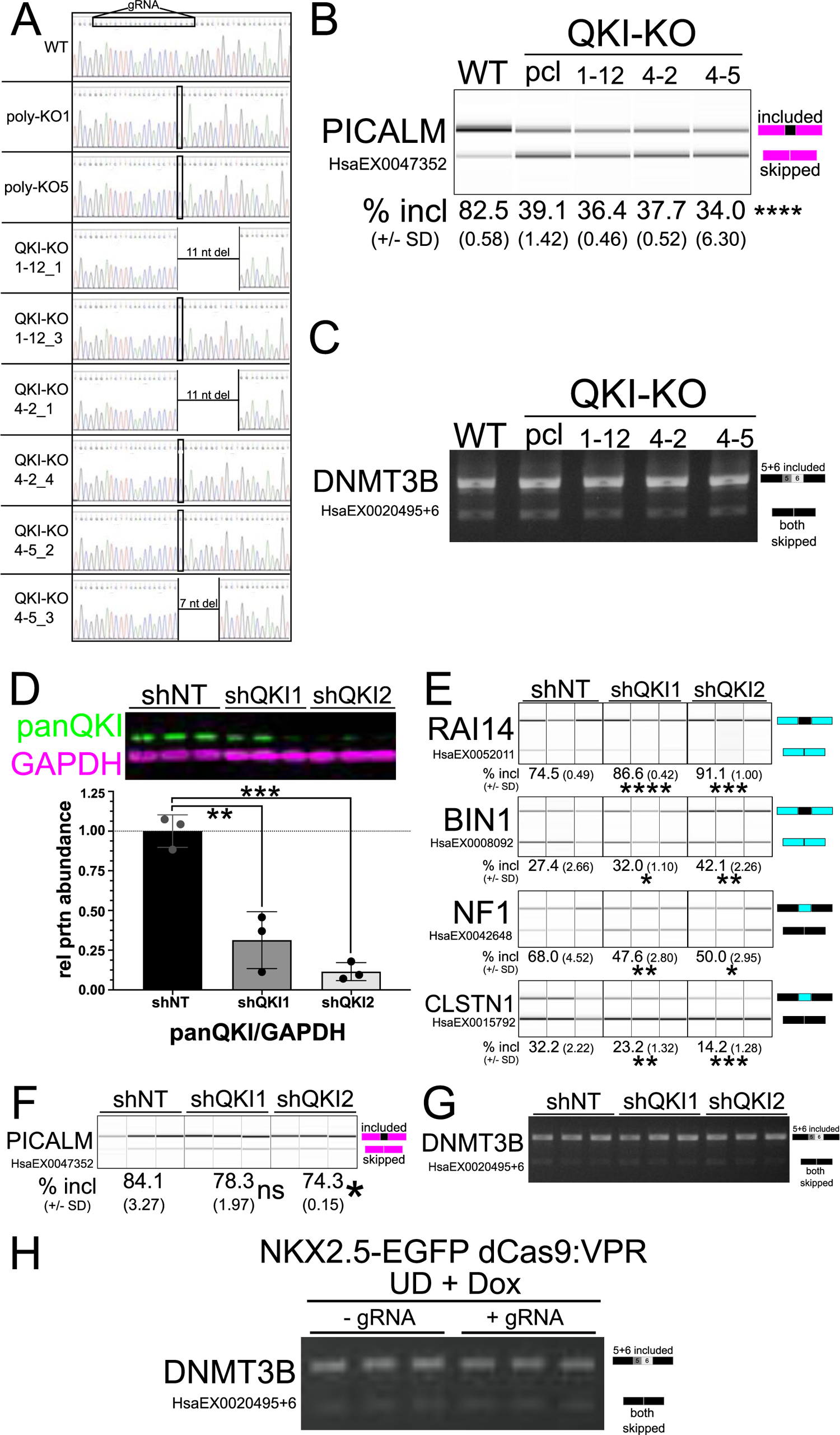
A. Chromatograms showing reads from genomic DNA amplified by PCR then cloned and sequenced from NKX2.5èGFP WT and QKI KO hESCs: the top panel shows WT sequence with guide RNA target noted in box. The next two panels show two independent sequences derived from polyclonal QKI KO cells (vertical box indicates nucleotide insertion); the following two panels show two independent sequences from single cell clone number 1-12 (deletion noted and vertical box indicates nucleotide insertion); the next two panels show two independent sequences from clone number 4-2 which has the same mutations described in clone 1-12; the final two panels show two independent sequences from clone number 4-5 which has a single nucleotide insertion (vertical box) and 7 nucleotide deletion. B. RT-PCR and BioAnalyzer gel-like image of RNA extracted from UD *NKX2.5*èEGFP hESCs described (see Figs 3A and 3B) above to measure alternative exon inclusion for *PICALM* (result shown is representative of 3 biological replicates with mean percent included +/- standard deviation shown below; each KO value is \*\*\*\**P* < 0.0001 by Student’s t-test). C. RT-PCR and agarose gel analysis of RNA extracted from UD *NKX2.5*èEGFP hESCs as described above to measure exon inclusion for *DNMT3B* (result shown is representative of 3 biological replicates). D. Western blot of 3 independent biological replicates of protein extracted from UD H9 hESCs, transduced with shNT, shQKI1, and shQKI2 probed with antibodies for panQKI (green) and GAPDH (magenta), and showing panQKI protein abundance relative to GAPDH in bar graph below, normalized to shNT (\*\**P* ≤ 0.01 and \*\*\**P* ≤ 0.001 by Student’s t-test). E. RT-PCR and BioAnalyzer gel-like image of RNA extracted from UD H9 hESCs transduced with shRNAs described above (in biological triplicate) to measure alternative exon inclusion of *RAI14*, *BIN1*, *CLSTN1*, and *NF1* (the mean values +/- standard deviation are reported below; \**P* < 0.05, \*\**P* < 0.01, \*\*\**P* < 0.001, \*\*\*\**P* < 0.0001 by Student’s t-test). F. RT-PCR and BioAnalyzer gel-like image of RNA extracted from UD H9 hESCs transduced in triplicate with shRNAs described above (see Fig 3E) to measure alternative exon inclusion of *PICALM* (the mean values +/- standard deviation are reported below; ns denotes not significant, \**P* < 0.05). G. RT-PCR and agarose gel analysis of RNA extracted from UD H9 hESCs transduced in triplicate with shRNAs described above to measure exon inclusion of *DNMT3B*. H. RT-PCR and agarose gel of RNA extracted from UD *NKX2.5*èEGFP dCas9:VPR hESCs (described in Figs 3C and 3D) to measure exon inclusion for *DNMT3B*.

**SUPPLEMENTAL FIGURE 4:**
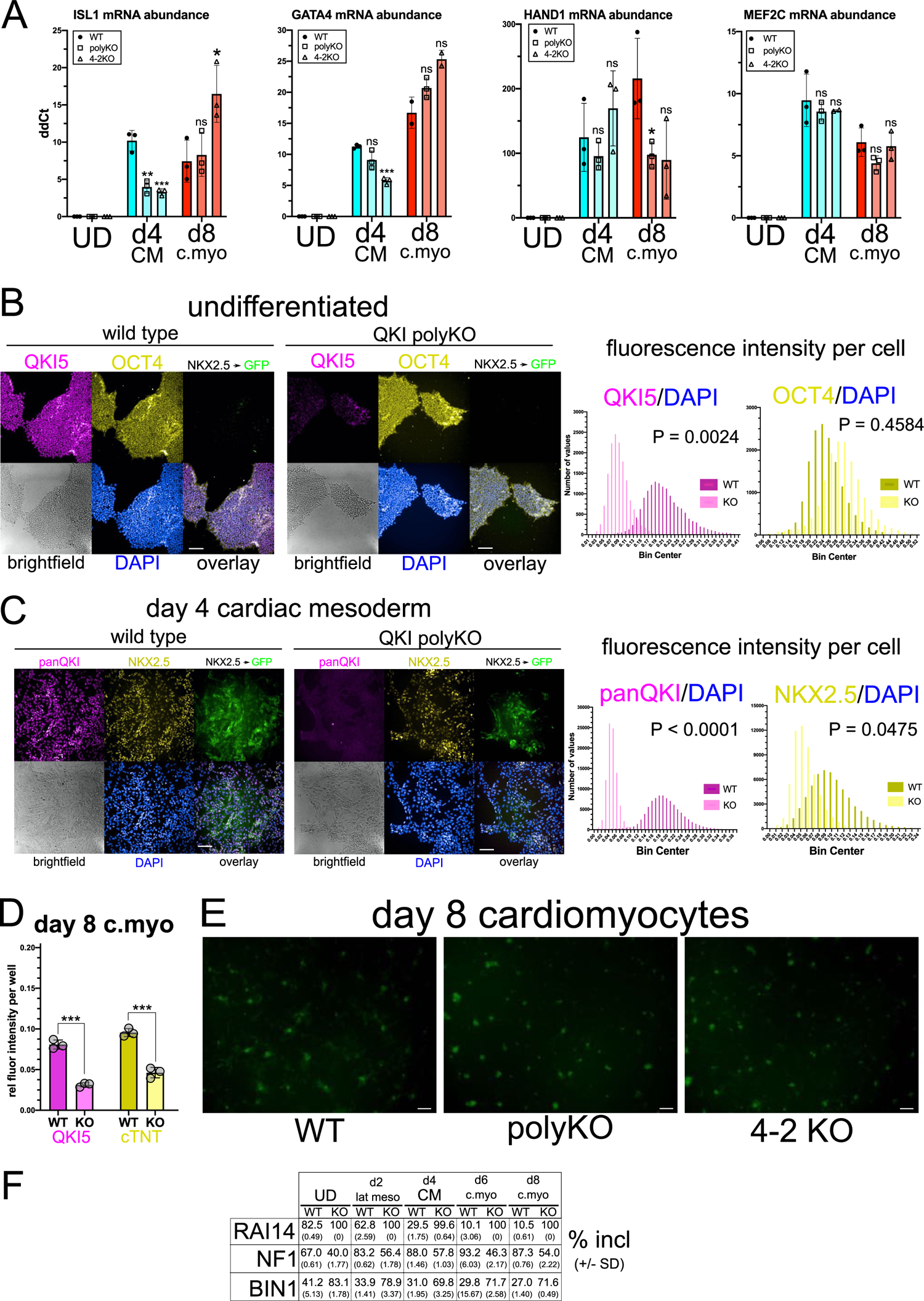
A. RT-qPCR measuring abundance of RNA extracted from *NKX2.5*èEGFP UD hESCs, d4 CM cells, and d8 c.myo cells for *ISL1*, *GATA4*, *HAND1*, and *MEF2C* mRNAs relative to *HMBS* mRNA in WT, polyKO, or 4-2KO cells (biological replicate n = 2 or 3 each; ns = not significant, \**P* ≤ 0.05, \*\**P* ≤ 0.01, \*\*\**P* ≤ 0.001 measured by Student’s t-test). B. Indirect immunofluorescence of UD *NKX2.5*èGFP WT (left) or polyclonal *QKI*-KO (right) hESCs analyzed by high content imaging, showing QKI5 (magenta), OCT4 (yellow), EGFP (green), brightfield (gray), and DAPI (blue) with each fluorescent channel overlaid. On the right, the fluorescence intensity values per cell (relative to DAPI) are shown for QKI5 and OCT4 with *P* values calculated by Mann-Whitney U. C. Indirect immunofluorescence of *NKX2.5*èGFP WT (left) or polyclonal *QKI*-KO (right) d4 CM cells analyzed by high content imaging, showing panQKI (magenta), endogenous NKX2.5 (yellow), EGFP (green), brightfield (gray), and DAPI (blue) with each fluorescent channel overlaid. On the right, the fluorescent intensity values per cell (relative to DAPI) are shown for panQKI and NKX2.5 with *P* values calculated by Mann-Whitney U. D. Fluorescent intensity values per well measured by indirect immunofluorescence as described in Fig 4F for QKI5 and cTNT (TNNT2), with each normalized to DAPI, in WT and polyclonal QKI-KO day 8 c.myo *NKX2.5*èGFP cells (n = 3 independent biological replicates; \*\*\**P* ≤ 0.001 by Student’s t-test). E. Low magnification epifluorescence imaging with FITC filter of live d8 c.myo *NKX2.5*èEGFP cells WT, polyKO, and 4-2KO (scale bar denotes 200 µm). F. Mean percent included values for alternatively spliced exons in *RAI14*, *NF1*, and *BIN1* calculated from three independent biological replicates, +/- standard deviation (see Fig 4H).

**SUPPLEMENTAL FIGURE 5:**
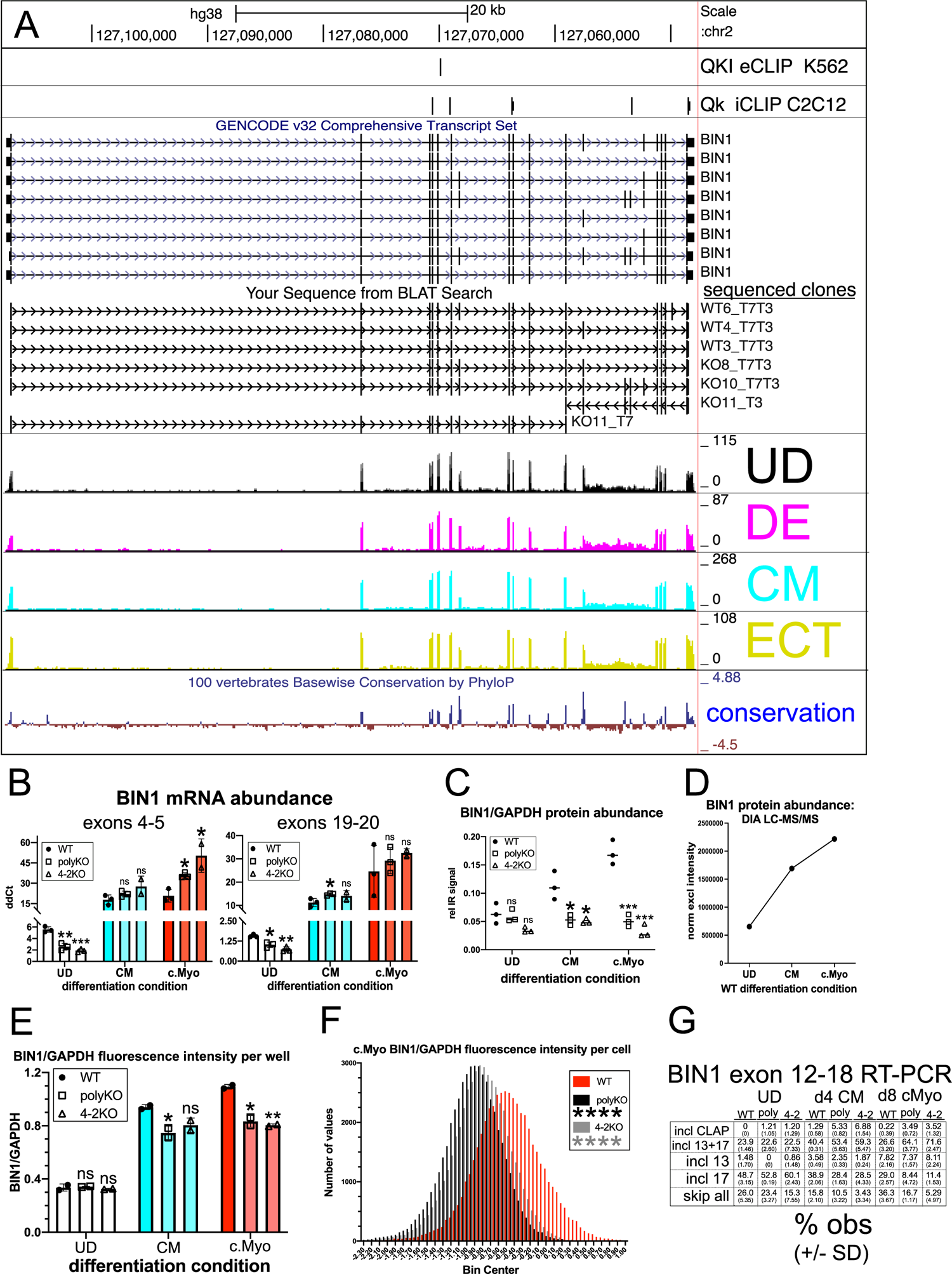
A. UCSC Genome Browser screen shot showing QKI eCLIP peaks from K562 cells, Qk iCLIP peaks from C2C12 myoblasts, Gencode Genes with annotated *BIN1* transcripts, unique full-length *BIN1* cDNA clones sequenced (Sanger sequencing) from UD and d8 c.myo WT and *QKI*-KO *NKX2.5*èEGFP cells (KO11 represents one full-length clone from two different forward and reverse sequencing reactions), coverage tracks from UD hESCs, DE cells, CM cells, and ECT cells RNA-seq datasets, and conservation tracks. B. RT-qPCR of RNA extracted from *NKX2.5*èEGFP WT, polyKO, or 4-2KO UD hESCs, d4CM cells, or d8 c.myo cells, and measuring either the 5’ end of *BIN1* (primer pair spanning exons 4 and 5) or the 3’ end of *BIN1* (primer pair spanning exons 19 and 20); value shown is ddCt relative to *HMBS*; n = 2 or 3 independent replicates, ns = not significant, \**P* < 0.05, \*\**P* < 0.01, \*\*\**P* < 0.001 by Student’s t-test compared to WT. C. BIN1 infrared protein signal relative to GAPDH infrared protein signal from western blots shown in Figs 4C, 4G, and 4K (n = 3 independent replicates, ns = not significant, \**P* < 0.05, \*\*\**P* < 0.001 by Student’s t-test compared to WT). D. BIN1 protein abundance (as normalized exclusive intensity on y-axis) from WT UD hESC, WT d4 CM, and WT d8 c.Myo cell extracts measured by data-independent acquisition LC-MS/MS. E. BIN1 fluorescence intensity relative to GAPDH per well (n = 2), measured by high content imaging in UD, d4 CM, and d8 c.myo *NKX2.5*èEGFP WT, polyKO, or 4-2KO cells (ns = not significant, \**P* < 0.05, \*\**P* < 0.01 by Student’s t-test compared to WT). F. BIN1 fluorescence intensity relative to GAPDH per cell (n = ∼50,000 cells, per cell type) in d8 c.myo *NKX2.5*èEGFP WT, polyKO, or 4-2KO cells (\*\*\*\**P* < 0.0001 by Mann Whitney U).

**SUPPLEMENTAL FIGURE 6:**
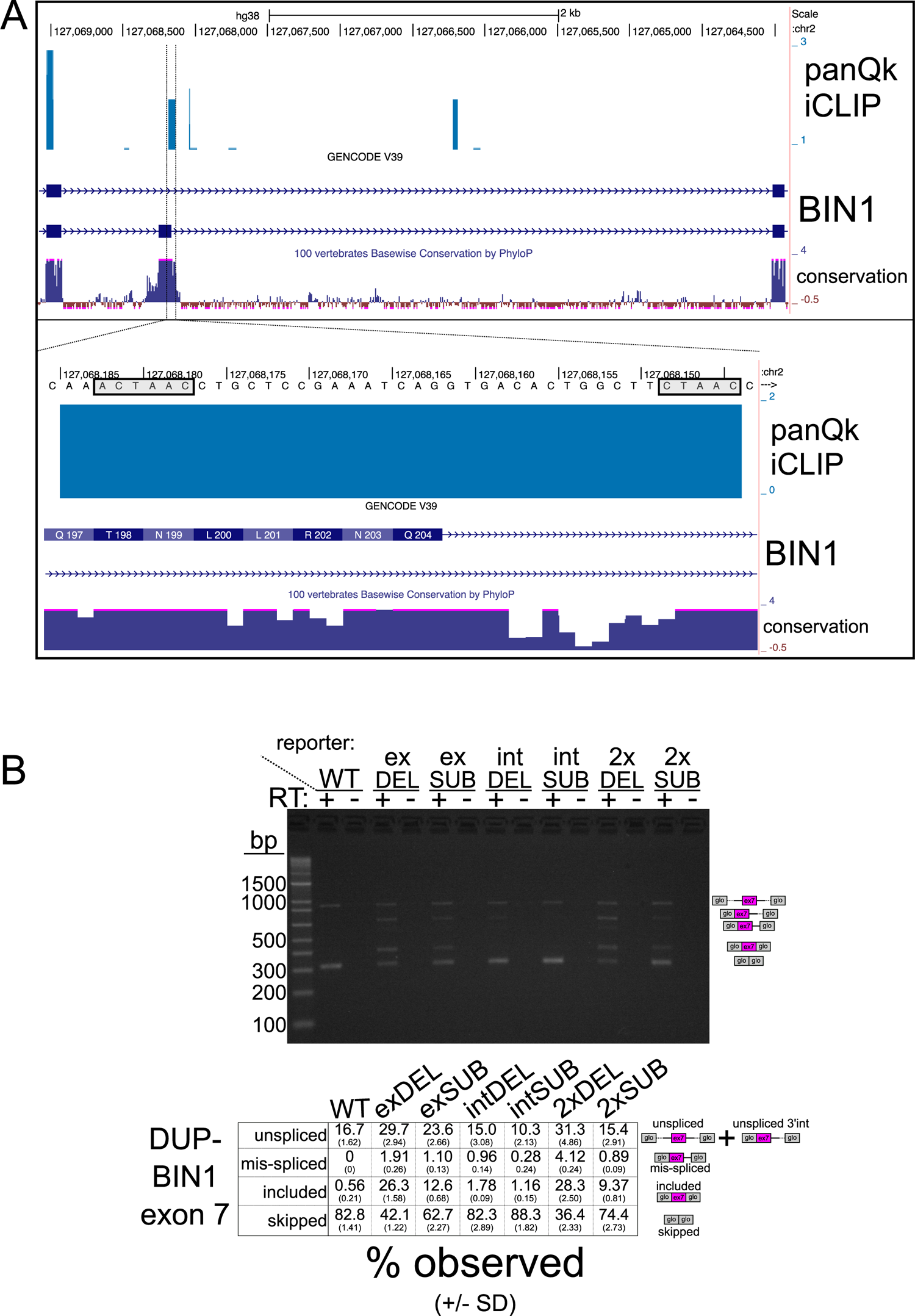
A. UCSC Genome Browser screen shot spanning BIN1 exons 6 through 8 (top) or zoomed-in (bottom) at the 3’end of BIN1 alternatively spliced exon 7, showing panQk iCLIP reads from C2C12 myoblasts (top), Gencode gene annotation (middle), and PhyloP conservation tracks (bottom). The lower inset shows genomic DNA sequences with boxes around the exonic ACTAAC and intronic CTAAC sequences of interest. B. Agarose gel showing PCR amplification of the samples used in Fig 6C either with (+) or without (-) reverse transcription (top), and mean values (+/- standard deviation) of PCR products shown in Fig 6C obtained by RT-PCR and BioAnalyzer quantification (bottom).

